# Single-residue effects on the behavior of a nascent polypeptide chain inside the ribosome exit tunnel

**DOI:** 10.1101/2024.08.20.608737

**Authors:** Fátima Pardo-Avila, Renuka Kudva, Michael Levitt, Gunnar von Heijne

**Affiliations:** Department of Structural Biology, Stanford University, Palo Alto, CA, USA; Department of Biochemistry and Biophysics, Stockholm University, SE-106 91 Stockholm, Sweden; Science for Life Laboratory Stockholm University, Box 1031, SE-171 21 Solna, Sweden

**Keywords:** Translational arrest, ribosome, SecM, Force Profile Analysis, molecular dynamics

## Abstract

Nascent polypeptide chains (NCs) are extruded from the ribosome through an exit tunnel (ET) traversing the large ribosomal subunit. The ET’s irregular and chemically complex wall allows for various NC-ET interactions. Translational arrest peptides (APs) bind in the ET to induce translational arrest, a property that can be exploited to study NC-ET interactions by Force Profile Analysis (FPA). We employed FPA and molecular dynamics (MD) simulations to investigate how individual residues within a glycine-serine repeat segment of an AP-stalled NC interact with the ET to exert a pulling force on the AP and release stalling. Our results indicate that large and hydrophobic residues generate a pulling force on the AP when placed ≳10 residues away from the peptidyl transfer center (PTC). An asparagine placed 12 residues from the PTC appears to form a specific stabilizing interaction with the tip of ribosomal protein uL22 that reduces the pulling force on the AP, whereas a lysine or leucine residue in the same position increases the pulling force. Finally, the MD simulations suggest how the *Mannheimia succiniciproducens* SecM AP interacts with the ET to promote translational stalling.

**Graphical Abstract:** 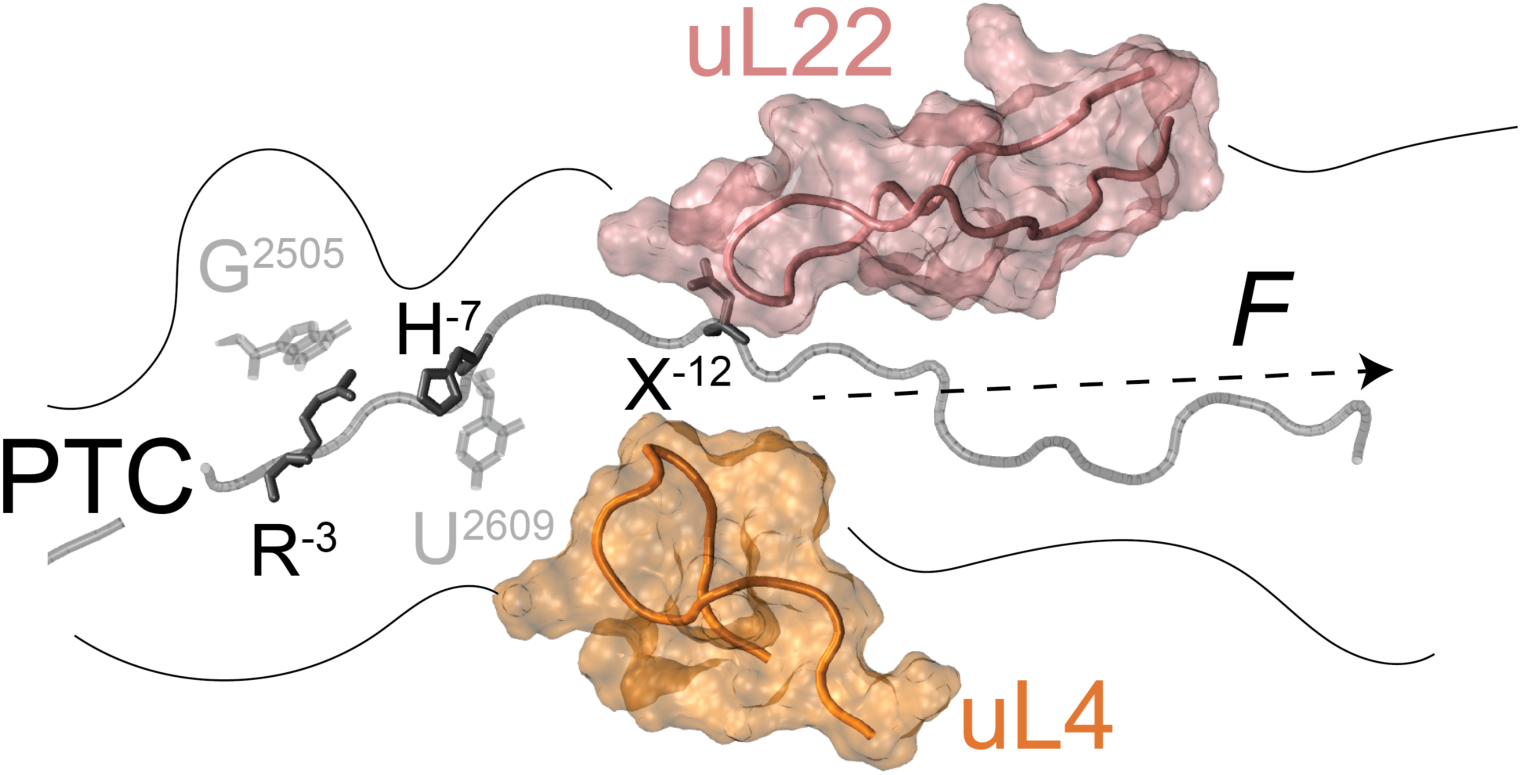

## Introduction

During translation, the elongating polypeptide chain traverses the ribosome through a ∼100 Å long exit tunnel (ET) in the large ribosomal subunit (1–4). While the ET was initially thought to have a “Teflon-like” surface that would have minimal interactions with the nascent polypeptide chain (NC) (3), it is by now well-established that there are ample possibilities for such interactions in specific regions along its length (5–14). The best-studied examples where interactions between the ET and the NC modulate translation elongation are the so-called translational arrest peptides (APs), relatively short stretches of sequences in NCs that interact with the proximal parts of the ET to induce translational arrest (15,16). Detailed information on AP-ET interactions has been obtained by cryo-EM and molecular dynamics (MD) simulations of ribosome-nascent chain complexes (RNCs) and by extensive mutagenesis studies (12–14,17–24), but studies of how individual residues may engage in NC-ET interactions in more distal parts of the exit tunnel are less common (25–29).

Here, we use Force Profile Analysis (FPA), a method based on the observation that the translational arrest efficiency of an AP is sensitive to external pulling forces acting on the NC, to map residue-specific interactions between individual amino acids introduced within a 19-residue segment comprising alternating glycine (G) and serine (S) residues that is stalled by the eight-residue long SecM AP from *Mannheimia succiniproducens* (SecM(*Ms*))(8) and the ET. Along with all-atom MD simulations, we examine the interactions of single charged (K, D), polar (N), and hydrophobic (W, P, L) residues at different locations in the GS-segment, corresponding to a part of the ET that is ∼20 Å to ∼70 Å distant from the peptidyl transferase center (PTC) of the ribosome. Overall, our results indicate that the introduction of larger residues in this area of the NC generates a pulling force on the AP in a direction away from the PTC. Moreover, placing an L or D residue in the nascent chain at position -18 (relative to the PTC), just beyond the constriction site formed by the loops of ribosomal proteins uL4 and uL22 protruding into the ET (30), leads to a specific increase in the pulling force. In contrast, an N residue placed at position -12 relative to the PTC (i.e., at the constriction site) appears to make a specific stabilizing interaction with the tip of protein uL22 that potentially reduces the pulling force on the AP, while a K or L residue in the same position increases the pulling force. In addition, data from MD simulations suggest how residues in the SecM(*Ms*) AP can interact with rRNA within the ET, potentially affecting translational stalling.

## Materials and methods

### Enzymes and chemicals

All enzymes used in this study were purchased from Thermo Fisher Scientific and New England Biolabs. T7-RNA polymerase was purchased from Promega. All reagents and chemicals except for Bacto-yeast extract and Bacto-peptone were purchased from Sigma-Aldrich; Bacto-yeast extract and Bacto-peptone to culture *E.coli* cells for the S30 extract, were purchased from BD-Biosciences. Oligonucleotides were from Eurofins Genomics and gene fragments from Thermo Fisher Scientific (GeneArt). DNA isolation/purification kits and precast polyacrylamide gels were from Thermo Fisher Scientific. L-[^35^S]-methionine was obtained from PerkinElmer.

### Cloning and Mutagenesis

The ADR1a constructs that were sub-cloned into the pET19b expression plasmid in a previous study (31) were used as a parental construct for this project. Gene fragments with DNA regions encoding for a total of 40 residues (consisting of alternating glycine and serine GS repeats) and the SecM(*Ms*) AP were designed and ordered from GeneArt (Thermo Fisher Scientific) – the codons encoding G and S were varied to avoid repeats. This fragment was introduced at the 3’ end of the gene encoding for ADR1a by Gibson assembly®. The shorter constructs for the initial screen to select the appropriate length of the GS-fragment for further experiments (ADR1 linker length scan available in Supplementary Excel file on Zenodo: 10.5281/zenodo.18988399) were generated by truncating the GS-segment to the desired length by inverse PCR using phosphorylated primers, ligation with T4 DNA ligase and transformation into DH5α chemical competent cells. The different single mutants (single amino acid substitutions) were generated from the construct with a 19 amino acid GS-segment and the eight amino acid SecM(*Ms*) AP (as described in the Results) by site directed mutagenesis using partially overlapping primers. All cloning and mutagenesis products were confirmed by DNA sequencing.

The amino acid sequences for all the constructs can be found in the Supplementary files on Zenodo (10.5281/zenodo.18988399).

### Coupled in vitro transcription and translation and quantitation

The constructs used in this study were translated in an *E.coli* Zn-free S30 cell extract (31). The S30 extract was prepared from *E.coli* MRE 600 cells following the protocol detailed in (32), with some modifications to rid the lysate of endogenous Zn^2+.^ Specifically, cells were cultured to an A600 of 1.2, following which they were treated with 100 µM TPEN for one hour, and harvested by centrifugation and prepared as described (32).

300 ng of plasmid DNA of the respective constructs were used as templates for polypeptide synthesis in the Zn-free S30 cell extract, and translation was carried out in the presence of [^35^S] Methionine at 37°C for 15 min and shaking at 300 r.p.m. (33). Since the S30 extract used was depleted of endogenous Zn^2+^, for the reactions + Zn^2+^, the reactions included 50 μM zinc acetate. The translation reaction was terminated after 15 min by treating the samples with a final concentration of 5% trichloroacetic acid (TCA), followed by a 30 min incubation on ice. They were subsequently centrifuged at 20,000 g for 10 min in a tabletop centrifuge (Eppendorf) and the pellet obtained was solubilized in Laemmli buffer, supplemented with RNaseA (400 μg/ml), and denatured at 37°C for 15 min.

The samples were resolved on 12% Bis-Tris gels (Thermo Scientific) in MOPS buffer. Gels were dried and subjected to autoradiography and scanned using the Fujifilm FLA-9000 phosphorimager for visualization of radioactively labeled translated proteins. The one-dimensional intensity profiles corresponding to the protein bands on each individual lane in the gel images were extracted using ImageGauge (Fujifilm). The output .txt files (deposited on Zenodo: 10.5281/zenodo.13244596) were used as inputs to visualize and fit to a Gaussian distribution using EasyQuant (Rickard Hedman, Stockholm University). The sum of the arrested and full-length bands was calculated, and this was used to estimate the fraction full-length protein for each construct as exemplified in Figure 1.

**Figure 1.**
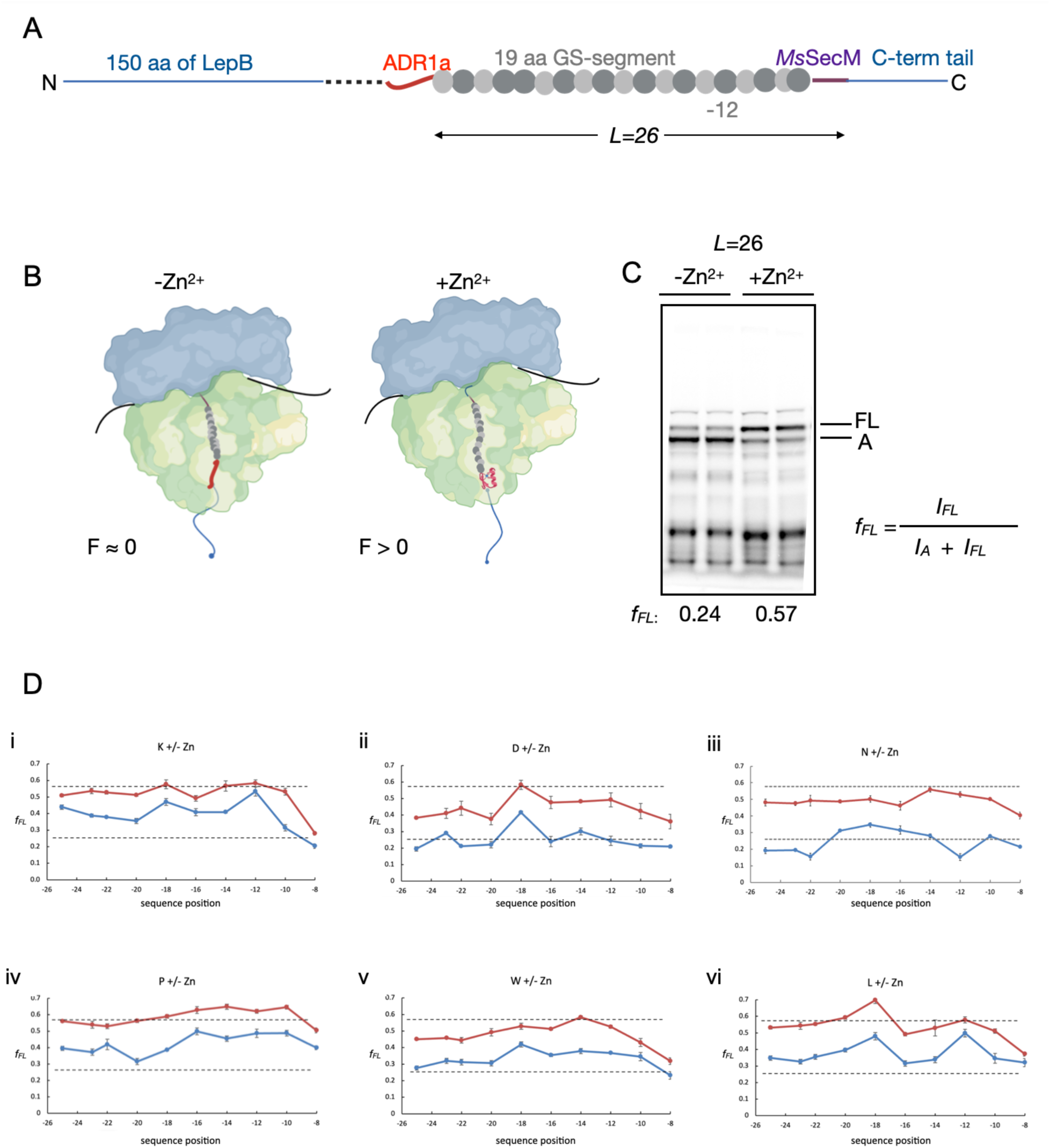
(A)Diagrammatic representation of constructs used. The Zinc-finger domain of ADR1a was fused to the SecM AP from *Mannheimia succiniciproducens* via 19 amino acid GS-repeats (GS-segment). 150 amino acids of the periplasmic domain of LepB were introduced at the N-terminus of ADR1 and 23 amino acids at the C-terminus of the SecM(*Ms*) AP to be able to resolve the arrested and full-length protein products by SDS-PAGE and autoradiography. (B) A schematic of ribosomes stalled by the SecM AP and how an N-terminal pulling force generated by the folding of ADR1a can result in a resumption of translation. In the left-hand panel, ADR1a (red) does not fold due to the absence of Zn^2+^ whereas ADR1a folds inside the ET in the presence of Zn^2+^, generating a pulling force on the NC (right-hand panel). (C) Autoradiographs of unfolded (-Zn^2+^) and folded (+Zn^2+^) ADR1a constructs stalled by the SecM(*Ms*) AP after radioactive pulse-labelling *in vitro* and SDS-PAGE. The relative amounts of arrested (*A*) and full-length (*FL*) product were estimated by quantification of the protein bands in the autoradiographs, and the fraction full-length was calculated as *f_FL_=I_FL_/(I_A_+I_FL_)*. Two repeat experiments are shown for each construct. (D) Force profile analysis of constructs with (i) K; (ii) D; (iii) N; (iv) P; (v) W, and (vi) L at different positions within the 19 residue Gly-Ser segment. Sequence positions are counted as in Fig. 2B, i.e., from the penultimate Ser residue in the SecM(*Ms*) AP.

### Model building

Our initial ribosome-AP model was based on the PDB structure *3jbu* (19). This structure is based on a cryo-electron density map of the *Escherichia coli* ribosome stalled during translation by the 17 amino acid long arrest peptide SecM. The nascent peptide chain modelled in *3jbu* contains 4 peptide bonds with the omega dihedral in the *cis* configuration (the cis peptide bonds in the *3jbu* nascent peptide are shown in bold: K^-24^LISE**ED**LFSTP**VW**ISQAQ**GI-RA**G^-1^, numbering according the SecM(*Ms*) NC) (SI Fig. SI2 and Fig. 2). It is highly unlikely that omega dihedrals adopt in the *cis* configuration, so we changed all these bonds to the *trans* configuration. We first used PyMOL (34) to mutate the residues in the nascent chain to match the sequence used in the experimental setup: the SecM(*Ms*) AP (with the penultimate S residue in the P-site), plus a 19-residue GS-segment in the G^-12^ model, and the N and K mutations at position -12 for the mutated models. (Fig. 2). Then, for the residues at the positions corresponding to E^-19^D^-18^ and R^-3^A^-2^, it was possible to fix the peptide bonds by “flipping” the oxygen of the peptide bond (manually change the coordinates of the atom), followed by energy minimization to correct the geometry. To fix the residues at the positions corresponding to V^-12^W^-11^ and G^-5^I^-4^, we used the GeneralizedKIC mover (35) implemented in Rosetta (36,37). We defined the loops as 4 residues centered at the *cis* bond. We fixed one bond at a time in a sequential manner, starting with the V^-12^W^-11^ position. We generated 100 models for each position and selected the one with the combination of the lowest score and lowest r.m.s.d. to the NC in the crystal structure *3jbu*. We used the fixed model and PyMOL to create a control model comprising 26 GS residues, and without the SecM(*Ms*) AP.

**Figure 2.**
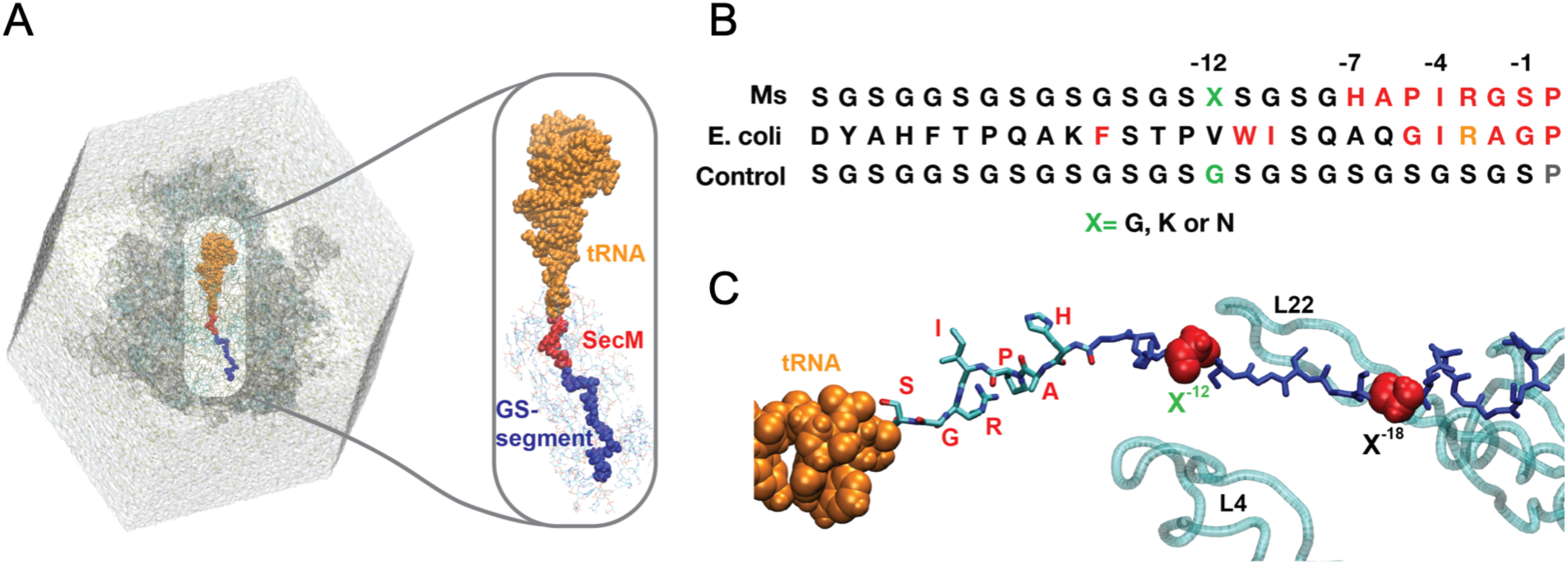
Molecular Dynamics setup. (A) Ribosome in a solvated dodecahedron box. The ribosome is shown as a cartoon, and the solvent as a grey surface. The exit tunnel is highlighted. The tRNA is shown as orange spheres, SecM(*Ms*) AP as red spheres, and GS-repeat linker as blue spheres. In the close-up, the nucleic acids within 15 Å of the nascent peptide are shown as lines. (B) SecM AP and GS-segment sequences. Residues critical for stalling in the SecM(*Ms*) and SecM(*Ec*) APs are highlighted in red. Position -12 is highlighted in green; X is either Gly, Lys, or Asn. The *E. coli* SecM(*Ec*) AP sequence is shown for reference. In the Control sequence, the SecM(*Ms*) AP is replaced by an equally long GS-repeat sequence. (C) Close-up of the nascent peptide (G^-12^ sequence) inside the exit tunnel. The loops of the uL4 and uL22 proteins are shown in cyan as references. Positions -12 and -18 are shown as red spheres.

To build the *8qoa* model (12), we obtained the coordinates from the PDB and used PyMOL to mutate the NC residues to match the SecM(*Ms*) AP (with the penultimate S residue in the P-site) and the 19-residue GS-segment.

### Building nascent peptide bound to tRNA

The nascent chain with the fixed peptide bonds was concatenated with the tRNA chain present in *3jbu*, and the residues were renumbered. A new bond had to be created between the Ser ^-1^ of the nascent peptide and the last nucleotide on the 3’ (RA104) of the tRNA. We obtained the missing parameters for the bond between the nucleic acid and the amino acid (Ser^-1^) using ACPYPE (38). The parameters necessary to specify the bond were added to the final topology files. The same protocol was followed for the *8qoa* model.

### Molecular Dynamics Simulations of the 3jbu systems

Molecular Dynamics simulations were performed using the Amber99SB-ILDN forcefield (39) with GROMACS 2018.4 (40). Each system was solvated in a dodecahedron box, with a minimum distance of 1.5 nm between the ribosome and the box edge. For all systems, the box vectors were 30.96 × 30.96 × 21.89 nm³. The systems were solvated with explicit TIP3P (41) water, and neutralized with K^+^. We then added Cl^-^ and Mg^2+^ ions to match a 10 mM concentration of MgCl_2_ to mimic the experimental conditions. In total, the unperturbed G^-12^contained 2,094,372 atoms, including 616878 water molecules, 4115 K^+^, 258 Cl^-^, and 129 Mg^2+^ ions. The Control system contained a total of 2,094,345 atoms, including 616,881 water molecules, 4116 K^+^, 258 Cl^-^, and 129 Mg^2+^ ions. The N^-12^ contained 2,094,358 atoms, including 616,871 water molecules,4115 K^+^, 258 Cl^-^, and 129 Mg^2+^ ions. The K^-12^ contained 2,094,371 atoms, including 616,873 water molecules, 4114 K^+^, 258 Cl^-^, and 129 Mg^2+^ ions.

We followed an equilibration protocol based on a previously described protocol for ribosome simulations (42):

0–5 ns: NVT and position restraints on all ribosomal heavy atoms (force constant of 1000 kJ mol^−1^nm^−2^).

5–10 ns: NVT, the position restraints force constant was linearly decreased to zero.

10–20 ns: NPT with a Berendsen barostat (43) with a coupling constant τp= 1ps and an isotropic compressibility of 4.5·10^−5^bar^−1^

After equilibration, for each system (Control, G^-12^, K^-12^, N^-12^), 5 simulations of 50 ns were performed in the NVT ensemble, with periodic boundary conditions. A 10 Å cut-off was used for van der Waals and short-range electrostatic interactions. The Particle-Mesh Ewald (PME) summation method was used for long-range electrostatic interactions (44). Verlet cut-off scheme was used (45). Covalent bonds were constrained using the LINCS algorithm (46). The integration time-step was 2 fs for all steps.

### Molecular Dynamics Simulations of the 8qoa system

Molecular Dynamics simulations were performed using the Amber99SB-ILDN forcefield (39) with GROMACS 2024.4 (40). The system was solvated in a dodecahedron box, with a minimum distance of 1.5 nm between the ribosome and the box edge. For all systems, the box vectors were 30.61 × 30.61 × 21.65 nm³. The systems were solvated with explicit TIP3P (41) water, and neutralized with K^+^. We then added Cl^-^ and Mg^2+^ ions to match a 10 mM concentration of MgCl_2_ to mimic the experimental conditions used in the FPA. The solvated system contained a total of 2,024,720 atoms, including 595,424 water molecules, 3944 K^+^, 248 Cl^-^, and 124 Mg^2+^ ions. We followed the same equilibration protocol as for the *3jbu* systems. After equilibration, 6 simulations of 275 ns were performed in the NVT ensemble under the same conditions as the MD simulations of the *3jbu* systems.

### Hydrogen bonds calculation

To facilitate the analysis, we filtered the MD trajectories to extract a subset of atoms. We extracted all atoms within a 15 Å radius of the NC backbone (the initial G^-12^ model structure after EM). This resulted in 334 residue trajectories. We further filtered this trajectory to extract the atoms closest to the PTC, resulting in a subsystem of 78 residues.

We calculated the prevalence of hydrogen bonds present in the 334 residue trajectories and in a subsystem of 78 residues near the PTC (for all systems). We used the baker_hubbard function (47) implemented in the MDTraj Python library (48). The function identifies hydrogen bonds based on cutoffs for the Donor – H…Acceptor distance and angle:

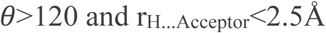

We used four frequency thresholds: 25%, 50%, 75%, and 90% of the simulation time.

### Cross-correlation analysis

The Pearson correlation of the covariance matrix (cross-correlation) allows us to quantify the similarity between two time-series datasets, by measuring the extent to which changes in one dataset corresponds to changes in another dataset, over a range of time. We can use cross-correlation to identify the coupling of the motion of the atoms in molecular dynamics simulations (49). Thus, being able to uncover hidden patterns or relationships. The correlation coefficient (C_ij_) between atom i and atom j is defined as:

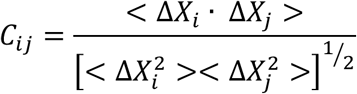

Where ΔX_i_ is the fluctuation of the position of atom i with respect to its mean position. C_ij_ = 1 denotes positively correlated motion, while C_ij_ = -1 denotes negatively correlated motion. We calculated the cross-correlation between the center of mass of all the residues in the 334-residue filtered trajectories. To calculate the cross-correlation, we first calculated the covariance matrix using the “gmx covar” function implemented in GROMACS (40), and then using the “cov2cor” function implemented in R 3.5.0 (50), to transform the covariance matrix to the cross-correlation matrix.

### Root Mean Square Deviation (RMSD) measurements (in layers)

The RMSD is the measure of the average distance between atoms of superimposed structures. We used pre-fitted center of mass (c.o.m.) trajectories, which were generated by calculating the c.o.m. of each residue in the structure for each frame of the trajectory. The initial model after energy minimization is the reference structure. To calculate the RMSD by layers, we divided the ribosome in 10 Å layers, using the nascent peptide as the center. All the layers were 10 Å, except for the last one, which included all atoms from 80 to 200 Å. We used the RMSD Trajectory Tool (RMSDTT v3.0 plugin) implemented in VMD (51) to both select the residues in each layer (all residues within X nm of NC and not within X-10 nm of NC; where X is goes from 10 to 80, except for the last layer, which included atoms from 80 to 100 Å) and calculate the RMSD to the initial conformation. For all trajectories of the same system, we used the same reference. The scripts used to plot the RMSD are deposited in Zenodo.

## Results

### Force Profile Analysis

FPA is based on the observation that the translational arrest efficiency of certain APs, including the SecM AP, is sensitive to external pulling forces acting on the NC (15,52,53). In general, stronger pulling forces result in low levels of translational arrest, and *vice versa*. Thus, by engineering an AP into a protein and measuring the arrest efficiency, one can obtain a proxy for the pulling force acting on the NC at the moment when the ribosome reaches the last codon in the AP (52). In FPA, a series of protein constructs are made where an AP followed by a short C-terminal tail is fused to a protein of interest, Fig. 1A. Each construct can either be translated *in vivo* or *in vitro* – in this study, we use an *in vitro* translation system consisting of a modified *E. coli* S30 cytosolic extract that is depleted of endogenous Zn^2+^ and devoid of membranes (32) – and subjected to a pulse of [^35^S]-Met, followed by analysis by SDS-PAGE and quantitation of the amount of product representing NCs arrested at the AP (*I_A_*) and the amount representing non-arrested, full-length chains (*I_FL_*), Fig. 1C. Finally, the fraction full-length product, *f_FL_* = *I_FL_*/(*I_A_*+*I_FL_*), is calculated as a measure of the pulling force exerted on the AP in that particular construct (52). FPA has been used to probe cotranslational processes such as protein folding (31,54–62), membrane protein biogenesis (52,63–65), and protein translocation across membranes (58,59,66). Here, we use FPA to probe NC-ET interactions by measuring how the pulling force exerted on the NC is modified by single point mutations introduced in different positions within a ribosome-embedded NC composed of glycine-serine (GS) repeats (referred to as the GS-segment).

The constructs have a common design that is based on previous studies on the cotranslational folding of the small Zn-finger domain ADR1a (31,62,67,68): a 150 residue long N-terminal unstructured segment derived from the periplasmic domain of the LepB protein followed by a 6-residue linker, the 29-residue ADR1a Zn-finger domain, a 19-residue GS-segment composed of 10 Gly and 9 Ser residues, the eight-residue *Mannheimia succiniciproducens* SecM (SecM(*Ms*)) AP (of sequence HAPIRGSP) – termed “linker or L” (see Figure 1A and C, and Supplementary Fig. S1A), and a 23-residue C-terminal tail. Zn^2+^-induced co-translational folding of ADR1a in the ET has previously been shown to generate a strong pulling force on the NC and to reduce the degree of translational stalling on the related *Escherichia coli* SecM AP (SecM(*Ec*)), while in the absence of Zn^2+^, ribosome stalling on the SecM(*Ec*) AP is efficient (31,62). The SecM(*Ms*) AP is the AP of choice in the present study since it is considerably shorter and more resilient to pulling forces than the SecM(*Ec*) AP (52), making it possible to probe locations in the NC closer to the PTC.

In order to validate the use of the SecM(*Ms*) AP in the S-30 *in vitro* translation system for this study, we recorded force profiles ±Zn^2+^ for ADR1a constructs with GS-segment lengths ranging from 11 to 31 residues; as shown in Supplementary Fig. S1A, the addition of Zn^2+^ resulted in a distinct increase in *f_FL_* values for linkers with 18-21 residues long GS-segments (refer to linker lengths 25-28 in Supplementary Fig. S1A), similar to what was found with the SecM(*Ec*) AP (31). We thus chose to use a 19-residue long GS-segment (corresponding to linker length 26 in Supplementary Fig. S1A) as a parental construct and as a reference for *f_FL_* in subsequent experiments.

To probe the effects on *f_FL_* of single charged (K, D), polar (N), and hydrophobic (W, L, P) residues in different positions along the NC, we analyzed six series of constructs, Fig. 1D, In each of the six scan-series, individual G residues in the 19-residue GS-segment were substituted by the specific tested residue type, and *f_FL_* was determined for each construct. For each of the six scan-series, we could thus determine the position-specific effect on *f_FL_* of the specific substituted residue in locations 8 to 25 residues (∼20 Å to ∼70 Å) away from the PTC, both in the presence (+Zn^2+^) and in the absence (-Zn^2+^) of a strong external pulling force on the NC.

In the six scan-series in Fig. 1D, *f_FL_* is plotted against the position of the mutated residue relative to the C-terminal Ser residue in the SecM(*Ms*) AP (which is attached to the P-site tRNA in the stalled ribosome-NC complex, counted as position -1). As expected, *f_FL_* values are always higher in the presence of Zn^2+^ (red curves) than in its absence (blue curves). Interestingly, the shapes of the *f_FL_* curves are similar ±Zn^2+^. However, in the presence of Zn^2+^, all *f_FL_* values are lower or equal to the *f_FL_* value for the unperturbed GS-segment (dotted line at *f_FL_* = 0.58), except for P in positions -10 to -16 and L in position -18. In the absence of Zn^2+^, *f_FL_* values are generally increased compared to the unperturbed GS-segment (dotted line at *f_FL_* = 0.26) for the K, W, L, and P series, but not for constructs in the D and N series. Finally, for all scans across the different series, *f_FL_* values drop in position -8, with the strongest drop seen in the scans recorded for the larger residues K, W, and L.

Comparing the different residue types, it is notable that the *f_FL_* values for the K series are higher than those for the D series, both in the presence and absence of Zn^2+^ (most easily seen in SI Fig. S1B). Although, at first sight, this might suggest that a positively charged residue exerts a stronger pulling force on the NC than a negatively charged one, this is inconsistent with the observation that the *f_FL_* values for the D and N series are similar throughout the range of positions tested (Supplementary Fig. S1C). Instead, this suggests correlation with the size of the residue: the series for larger and hydrophobic residues (K, W, L, P) generally have higher *f_FL_* values than those for the smaller ones (D, N). Overall, the P series has the highest *f_FL_* values of all the series in the region -8 to -16, i.e., in the proximal part of the ET, closer to the peptidyl transferase center (PTC) (Fig 1D).

Beyond these general trends, a few constructs have notably high or low *f_FL_* values in certain positions: D and L (and to some degree K and W) in position -18 (high *f_FL_*), L and K in position -12 (high *f_FL_*), and N in position -12 (low *f_FL_*). The latter result is particularly interesting, as it potentially suggests that an N residue at position -12 forms a specific interaction with the exit tunnel that increases the stalling strength of the AP, thereby reducing *f_FL_*.

### All-atom MD simulations

All-atom MD simulations were used to visualize and to help identify the molecular interactions that could result in the observed differences in *f_FL_* values obtained in the absence of Zn^2+^ (*i.e.*, under low pulling-force load) in the FPA experiments (Fig. 2A). We specifically focused on constructs with the G→N and G→K mutations in position -12 (referred to as N^-12^ and K^-12^, respectively) as these showed the largest observed difference in *f_FL_* values among the analyzed positions (Fig. 1D). The N^-12^ and K^-12^ mutants were compared to the unperturbed G^-12^ sequence, and to an additional control construct where the SecM(*Ms*) AP was substituted by an equally long stretch of eight residues consisting of GS-repeats (Fig. 2B).

Since there are currently no structures available for the *E. coli* ribosome with a stalled SecM(*Ms*) AP, and the best-resolved cryo-EM structure that was available for the SecM(*Ec*) AP during this study was *3jbu*, we generated a molecular model based on that structure (PDB ID *3jbu*; EMDB ID EMD-6483) (19). The amino acid residues in the NC for *3jbu* were substituted to match the sequence used in the experimental set-up described above: the SecM(*Ms*) AP (with the penultimate S residue in the P-site), plus a 19-residue GS-segment in the G^-12^ model, and the N and K mutations at position -12 for the mutated models. We also set-up a Control model consisting of 26 GS residues (Fig. 2B, C). Based on the experimental data that suggests a residue-specific effect from individual substituted amino acids within the NC on the pulling force that appears to be independent of the presence of Zn^2+^(and therefore the folding of ADR1a), we excluded ADR1a from the MD models.

The *3jbu* model that was used as a reference to build our SecM(*Ms*) AP model contained 4 peptide bonds with the omega dihedral in the *cis* configuration in the NC (peptide bonds shown in bold: KLISE**ED**LFSTP**VW**ISQAQ**GI**-**RA**G). All the *cis* peptide bonds were converted to the *trans* configuration, as it is unlikely that the NC would contain *cis* peptide bonds (69,70). Remodeling the backbone resulted in changes in sidechain orientation. In particular, R^-3^ no longer pointed towards the A-site (the A-site tRNA was not included in our model, SI Fig. S2).

While *3jbu* was the highest-resolution structure available for the SecM(*Ec*) AP when this study was set up and underway, in a recently published high-resolution ribosome structure with a stalled SecM(*Ec*) AP (12) (PDB ID *8qoa*; EMDB ID EMD-18534), the proximal SecM(*Ec*) region between F^150^-G^165^ forms a compact α-helix that is not present in the *3jbu* model. This α-helix was found to be stable during an extended MD simulation (12), and was also predicted by AlphaFold2 (71) and the secondary structure prediction program PSIPRED (72). Notably, neither AlphaFold2 nor PSIPRED predict a helical structure for the SecM(*Ms*) AP (SI Fig. S3). To test whether the helix observed in *8qoa* remains stable when its residues are mutated to match SecM(*Ms*), we performed MD simulations based on the *8qoa* model, using the backbone of the NC, but mutating the side chains to match the SecM(*Ms*) + GS-segment system (SI Fig. S4A). Six production runs, each 275 ns long were generated to track the secondary structure of the NC throughout the simulations. Consistent with the PSIPRED and AlphaFold2 predictions, the initial α-helix structure was rapidly lost in all six simulations (SI Fig. S4B-C and SI movies) even though no external pulling force was applied to the NC. This indicates that the SecM(*Ms*) AP is unlikely to adopt the compact conformation that is observed for the SecM(*Ec*) AP, and that our model of the SecM(*Ms*) + GS-segment (based on *3jbu*, with the *cis*-*trans* corrections noted above) therefore represents a reasonable starting point for the MD simulations.

Thus, we proceeded with the analysis of the simulations generated from the original models built from the *3jbu* structure. For each of the four systems (G^-12^, K^-12^, N^-12^, Control), the production runs consisted of five 50-ns replicates. We focused the analysis on the residues within a 15 Å radius of the NC (Fig. 2A).

No major differences in the ribosome structure between the replicas or between the four simulated systems were observed. When we calculated the root mean squared distance (RMSD) of the whole ribosome relative to the initial conformation, we obtained higher values than expected. From visual inspection of the simulations, we noticed that the L1 stalk of the large ribosomal subunit was very flexible. Thus, to verify that the ribosome core was stable, and that the high RMSD values we observed were due to the L1-stalk, we calculated the RMSD by dividing the ribosome into concentric atom layers, starting from the nascent peptide (SI Fig. S5). The RMSD plots show the similarity to a reference structure – the original ribosome model used to start all the simulations after energy minimization. As seen in the SI Fig. S5A, all the simulations were stable and behaved similarly.

We further investigated the stability of the residues of the ET by calculating the root mean squared fluctuation (RMSF), which measures how much a residue moves during the simulation (fluctuation around the average position) for the components of the exit tunnel (nucleic acids of the tunnel, P-site tRNA, the loops of the proteins uL22, uL4, and uL23 that protrude into the ET), as well as for the NC. The residues in the protein loops have low RMSF values, and the values are similar for all simulations (SI Fig. S6). We observed higher RMSF values for the nucleic acids in the ET than for the amino acids in the uL22 and uL24 loops; the values were again similar among all four systems. Notably, the RMSF values for the K^-12^ and N^-12^ systems are very similar along the NC, except in the region between the SecM(*Ms*) AP and the mutation site (residues -12 to -7), where the K^-12^ NC appears more mobile than the N^-12^ NC (SI Fig. S6A).

In short, the ET components behave similarly in all the simulations and are stable regardless of the specific sequence of the nascent peptide.

### NC-ET interactions

To investigate whether the K^-12^ and N^-12^ mutations might affect NC-ET interactions and therefore explain the observed experimental differences between the two sets of constructs, we calculated the fraction of the total simulation time that specific nucleotides or amino acid residues in the ET spend within a distance of 4 Å of residues S^-11^, X^-12^, and S^-13^ in the NC. The observed interactions involve a few distinct regions in the ET (Fig. 3). At position -12 in the NC, N^-12^ is the only residue tested that did not appear to interact at all with uL4. Both the K^-12^ and N^-12^ residues interact with uL22, A751, and A752 for more than 50% of the simulation time. While the interaction between N^-12^ and uL22 appears to be mainly due to sidechain-sidechain interactions, the interaction between K^-12^ and uL22 is through the K backbone atoms, with its sidechain pointing towards A751 and A752 (SI Fig. S7). The K^-12^ sidechain can also reach a pocket formed by A789/A790 and U1781.

**Figure 3.**
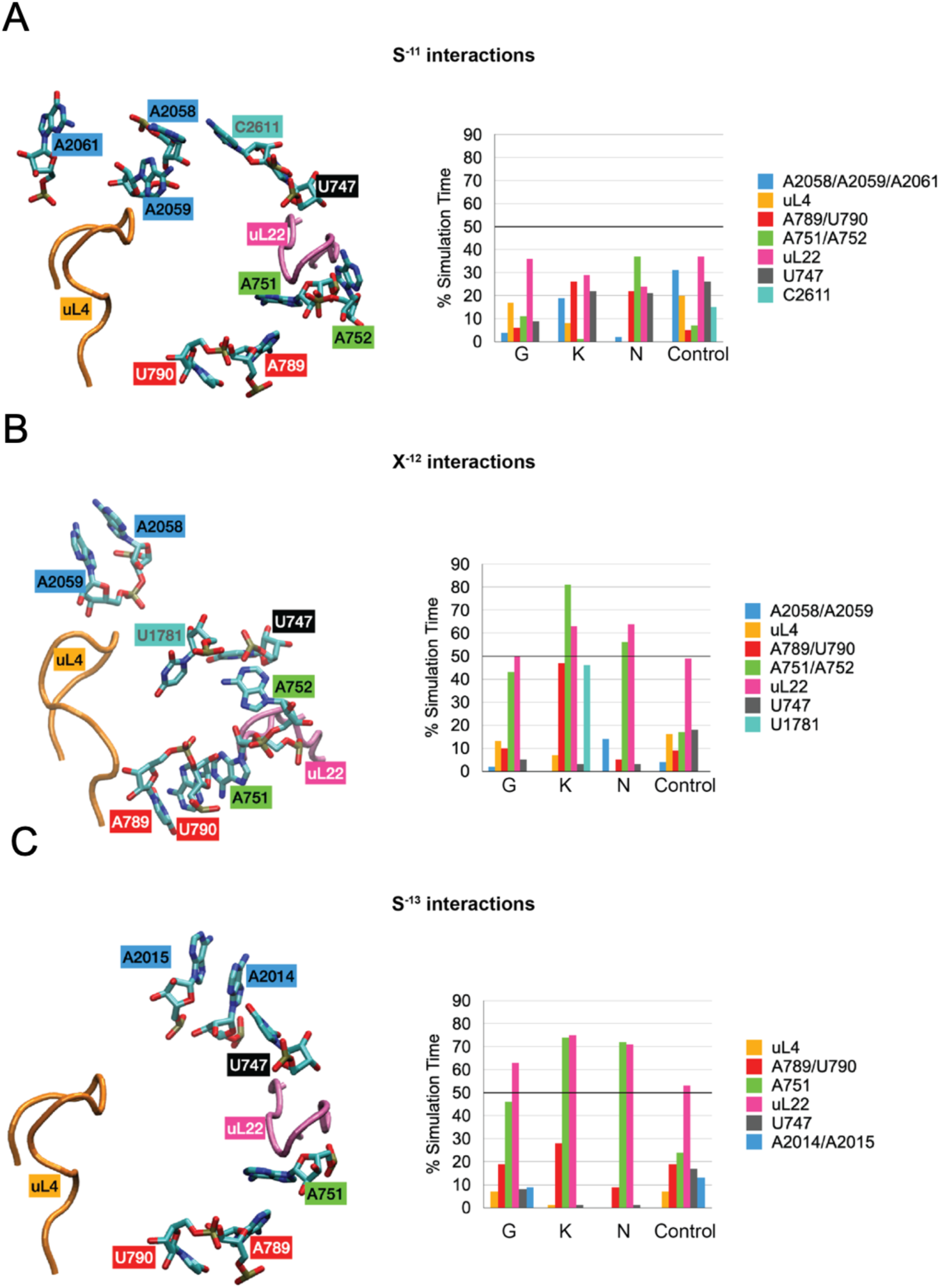
NC-ET interactions for residues S^-11^ (A), X^-12^ (B), and S^-13^ (C). Residues are defined as interacting when the minimum distance between them is less than 4 Å. The bar graphs show the percentage of the simulation time that the indicated residues interact. The interacting residues are shown in stick representations (left panels).

The interaction between S^-13^ and the exit tunnel follows a similar pattern, with the N^-12^ mutation strongly promoting interactions between S^-13^ and uL22/A751. In contrast, for S^-11^, we observed that consistent high-probability interactions are lacking, as most interactions are observed only in one single simulation (SI Fig. S8).

The N^-12^ mutation promotes a medium-probability interaction between S^-11^ and A751/A752, which is not seen in the other systems.

The 23S A751/A752 loop has been previously implicated in the ribosomal stalling in response to the SecM(*Ec*) and TnaC APs (73). Another study suggested the importance of stable aromatic interactions between W^155^ in the SecM(*Ec*) AP and A^751^ for ribosomal stalling (18).

Overall, we observe that N^-12^ and S^-13^ in the N^-12^ NC interact almost exclusively with uL22 and A751/A752. The N^-12^-uL22 interactions are mediated through the N^-12^ sidechain. In contrast, we observe that K^-12^ explores more extensive regions within the ET and interacts with uL22 via its backbone. These results are consistent with the higher RMSF values observed for the K^-12^ NC in the region around the mutation site (SI Fig. S6-S10).

### A persistent interaction between N^-12^ and uL22

To further characterize the interactions between the NC and the ET, we calculated the Pearson correlation of the covariance matrix of the center-of-mass of all residues (nucleotides and amino acids) within a 15 Å radius from the NC. We used the correlation to assess if the motion of a pair of residues is coordinated and to identify clusters of residues that could form a network of interactions that may affect translational arrest.

For the N^-12^ NC simulations, we found a strong correlation (cross-correlation values higher than 0.6) between N^-12^ and residues in the uL22 loop (Fig. 4A, SI Fig. S11 and S12). G^-12^ in the unperturbed NC variant also had a fairly strong correlation (cross-correlation values higher than 0.5) with the uL22 loop, whereas K^-12^ showed no such correlation. In contrast, none of the NC variants showed a strong correlation with the loop of uL4, which also forms part of the constriction site. Notably, mutations in Gly^91^ and Ala^93^ at the tip of the uL22 loop suppress the translational arrest induced by the SecM(*Ec*) AP (5). At the same time, uL4 has been suggested to be less involved in interactions that promote translational arrest by *E. coli* SecM (74), and mutations in uL4 had little or no effect on the SecM(*Ec*) AP response (75).

**Figure 4.**
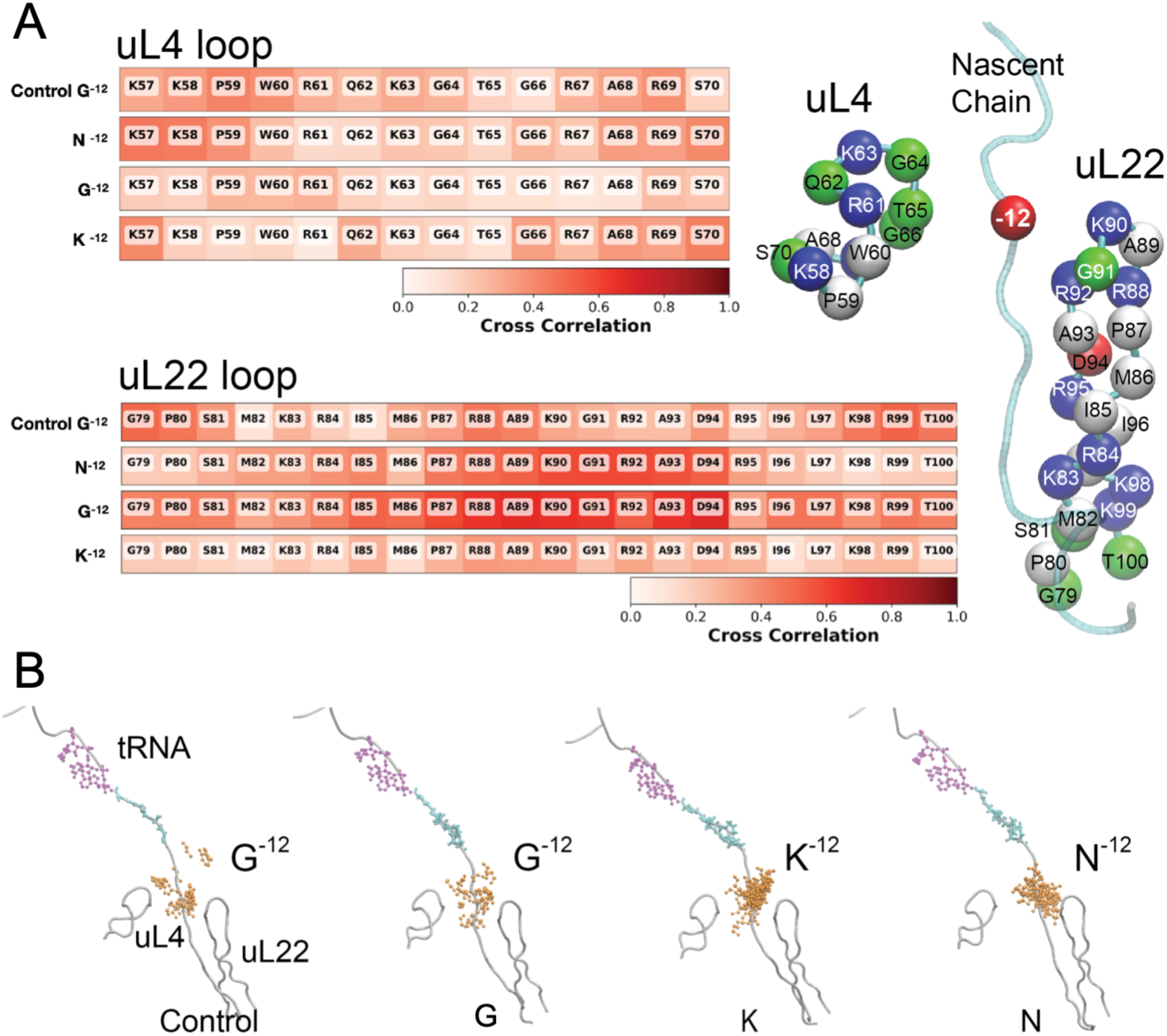
NC-ET interactions. (A) Cross-correlation between the residue of the nascent peptide in position -12 (for four systems) and the loops of proteins uL22 (residues G^79^ to T^100^) and uL4 (residues K^57^ to S^70^). The image on the right shows the loops of proteins uL22 (residues G^79^ to T^100^) and uL4 (residues K^57^ to S^70^). Each residue is represented as a sphere located at the center of mass of each residue. (B) The location within the ET of the nascent peptide residue at position -12. For each system, the average position of the loops of proteins uL4 and uL22 and the nascent peptide are shown in grey as cartoon representations. The SecM(*Ms*) AP residues are shown as cyan sticks, and the CCA terminus of the tRNA is shown as magenta sticks. To visualize the regions that the residue at position -12 visits during the simulation, this residue is shown as a CPK model (in orange) from overlapping frames of the sub-sampled trajectories.

Thus, we propose that the low *f_FL_* value recorded for the N^-12^ mutation (Fig. 1D) might result from a strong interaction between N^-12^ and uL22 that makes the NC bind more tightly to the ET and hence increases the arrest potency of the AP. While G^-12^ in the unperturbed system also has a strong cross-correlation with the uL22 loop, K^-12^ has a much weaker cross-correlation (values under 0.5) with the uL22 loop, in agreement with the higher *f_FL_* value of the latter.

### Hydrogen bonds and stacking interactions in the ET

In a final analysis (Fig. 5), we identified all the hydrogen bonds present within a 15 Å radius of the NC in the simulations. The hydrogen bonds that are present >50% of the simulation time are mostly between nucleotides. By lowering the threshold to 25% of the simulation time, we could identify a few hydrogen bond interactions between the NC and the ET in the region near the mutation site, including K^-12^ and N^-12^, that can both form a hydrogen bond with the backbone of K^90^ of uL22. As noted above, these interactions differ because the bond formed with K^-12^ is through its backbone, while N^-12^ interacts with uL22 via its sidechain.

**Figure 5.**
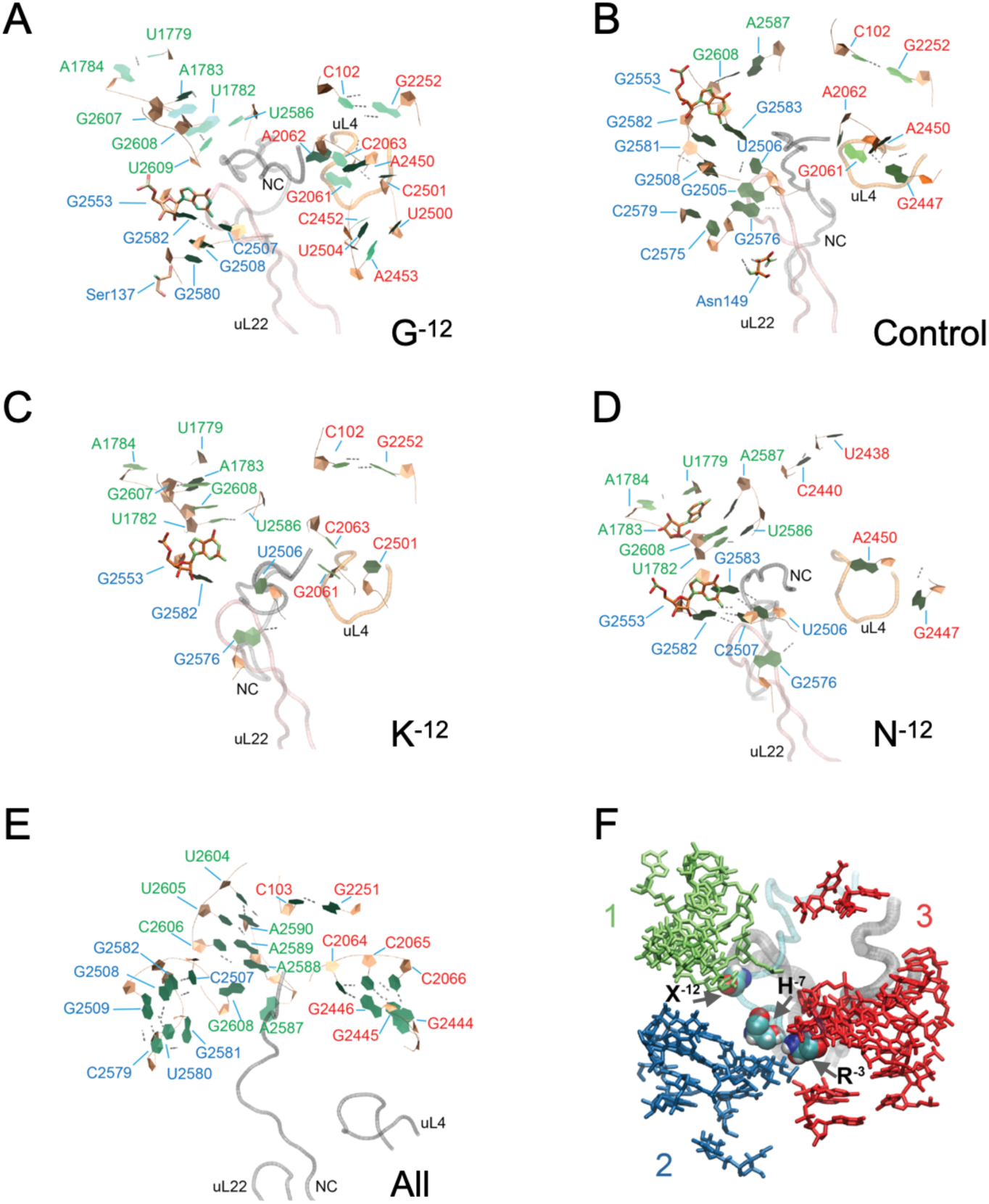
Hydrogen bond networks near the PTC. (A-E) The structures show the nucleic acids (orange and green, NewRibbons VMD representation) that form hydrogen bonds present at least 50% of the simulation time. Grey dashed lines between residues represent hydrogen bonds between residues. The loops of proteins uL4 (orange) and uL22 (pink), as well as the nascent chain (grey) are shown as tubes. Note that the frame selected for each system may show only some of all the possible hydrogen bonds. For a list of the hydrogen bonds present 50%, 75%, and 90% of the simulation time, refer to SI tables 1-3. To facilitate visualization, the nucleic acids that were involved in hydrogen bonds more than 50% of the time and were present in all the systems are shown on panel E and are not shown in panel A to D (A: G^-12^; B: Control; C: K^-12^; D: N^-12^). (F) The h-bonds are localized in three major regions, as shown in the blue, green, and red sticks. The loops of proteins uL22 and uL4, tRNA CCA fragment, and nascent peptide are shown as ribbons. SecM residues R^-3^, H^-7^, and X^-12^ are shown as spheres. The viewpoint is different from the one shown for panels A to E. The structure was rotated ∼90 degrees so the tRNA would come off the page towards the reader. The colors of the residue labels in panels A to E correspond to the three major regions highlighted in this panel.

We also identified hydrogen bonds near the PTC (Supplementary Tables 1 to 3). Three hydrogen bond clusters were found in this region, Fig. 5F. Cluster 2 is more extensive in the N^-12^ NC variant, including hydrogen bonds between U2506 and G2583, nucleotides that have been previously associated with stalling (76). Major differences can be observed between the SecM(*Ms*) AP simulations and the Control simulations (Fig. 5A-D). While in the SecM(*Ms*) AP simulations, R^-3^ in the SecM(*Ms*) AP and G2505 interact, either through hydrogen bonds or by stacking (SI Fig. S13), for the Control system hydrogen bonds are also formed between G2505 and G2581. Notably, in the SecM(*Ms*) AP simulations, rRNA bases U1782-U2586 and U1779-A1784 near the PTC interact, while these interactions are not seen in the Control system. U2586 has been previously proposed as relevant for SecM-mediated stalling (19). We also identified two regions that are stable for all four systems (low RMSF values and multiple hydrogen bonds): residues C2507-G2582 (near the A-site), and C2064-G2446, C2065-G2445, and C2066-G2444 (near the uL4 loop).

We further observed a stacking interaction between H^-7^ in the SecM(*Ms*) AP and U2609, present mainly in the N^-12^ NC variant, Fig. 6. Even though it only accounts for 20% of the frames, it potentially represents another stabilizing interaction between the N^-12^ NC and the ET. U2609 has been previously implicated in the ribosomal response to the SecM(*Ec*) AP (73), and the recent SecM(*Ec*) AP model *8qoa* shows a stacking interaction between the similarly placed F^150^ and U2609. In the TnaC structure (14,77), it was observed that U2609 is part of the binding pocket where the L-Trp molecule binds and has been found essential for stalling.

**Figure 6.**
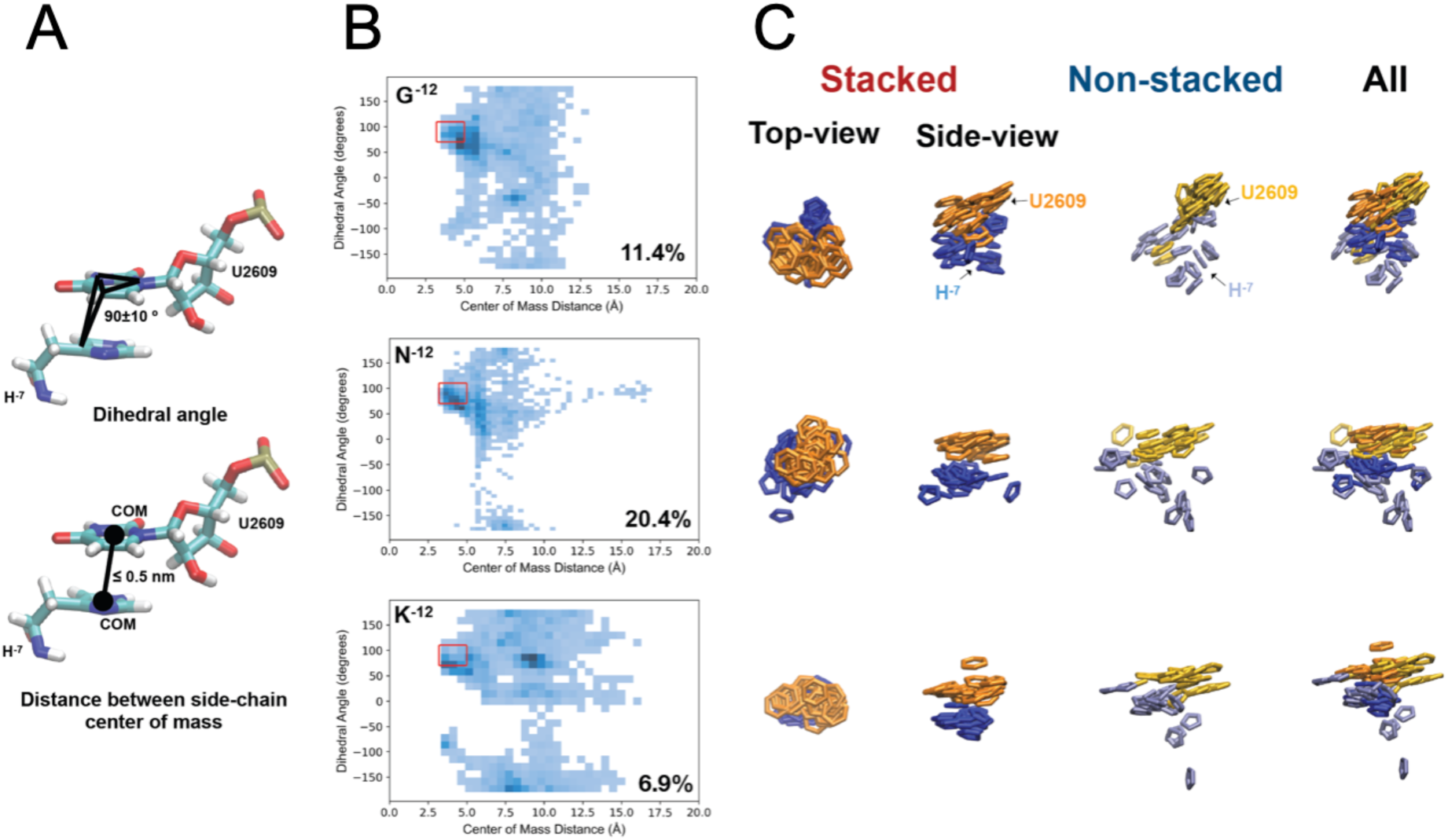
Stacking between H^-7^ and U2609. (A) Two reaction coordinates are used to define stacking between H^-7^ and U2609 (left panel). The distance between the center of mass (c.o.m.) of the H^-7^ sidechain and the base ring of U2609 is less than 0.5 nm, and the dihedral angle formed between the base ring of U2609 and the C2 atom of H^-7^ has a value between 70 and 110 degrees. All the frames were projected onto these two reaction coordinates (dihedral angle vs c.o.m. distance). (B) 2D histograms of the projections onto the reaction coordinates described in A. The red rectangle shows the region where the stacking conditions are met, and the percentage of frames that fall within this region is shown in the bottom right of each plot. (C) Extracted structures from the stacked and non-stacked regions highlighted in B. The aromatic rings of H^-7^ and U2609 are shown as blue and orange sticks, respectively.

## Discussion

In summary, under conditions of high external pulling force on the NC (in the presence of Zn^2+^), mutation of the G residues in the “skinny” GS-repeat sequence in positions -8 to -25 have no or a slightly reducing effect on *f_FL_*. Thus, not unexpectedly, the force generated by the cotranslational folding of ADR1a largely swamps out the effects of single point mutations in the GS-repeat sequence. In the absence of a strong external pulling force (no Zn^2+^), however, differences between the different types of residues start to appear. Substituting individual G residues in the GS-repeat segment to any of the larger residues (K, W, L) leads to increases in *f_FL_*; *i.e*., large residues tend to pull the NC towards the tunnel exit, away from the PTC. As the ET gets progressively wider beyond the constriction site, sidechain entropy would favor the movement of large residues in this direction, although we cannot exclude other explanations. Mutation of G residues to medium-size residues (D, N) does not, in general, affect *f_FL_* except in certain positions: -18 for D and N (high *f_FL_*) and -12 for N (low *f_FL_*). Previous experimental work (25,28,57) and theoretical calculations (78–82) have suggested that the average electrostatic potential varies along the ET; we do not see much evidence for an electrostatic effect (compare the *f_FL_* plots for the negatively charged D and its neutral analog N, Supplementary Fig. SI1B), implying that, on the single-residue scale, local residue-residue interactions (hydrogen bonds, salt bridges, stacking) dominate over electrostatic gradients.

Gersteuer et al. have proposed that the SecM(*Ec*) AP comprises two modules: an arrest module and a regulator module (12). The arrest module is formed by residues RAG/P (with the P residue attached to the A-site tRNA) and is directly involved in preventing peptide bond formation. Meanwhile, the more N-terminal regulator module can modulate the strength of stalling. In the SecM(*Ms*) AP, the arrest module would correspond to the RGS/P sequence. A mutagenesis scan showed that the SecM(*Ms*-Sup1) AP sequence is a stronger staller than SecM(*Ec*), as mutations that match SecM(*Ec*) (G^-2^→A and S^-1^→G) are significantly weaker stallers than SecM(*Ms*-Sup1). The mutagenesis scan also showed that R^-3^ is crucial for stalling (21).

U2504 has been proposed to control the access to the cavity where the incoming A-site amino acid would bind (83). In the old SecM(*Ec*) AP structure *3jbu* (19), the sidechain of R^-3^ (R^165^) also points towards this pocket, potentially blocking the A-site. However, in the recent, high-resolution structure of the SecM(*Ec*) AP *8qoa* (12), R^165^ forms a stacking interaction with U2504. Interestingly, even though we base our model of the SecM(*Ms*) AP on *3jbu*, our correction of a couple of presumably incorrectly modeled *cis* peptide bonds led to a starting structure for the MD simulations where the sidechain of R^-3^ no longer points to the same location as in *3jbu*. Instead, it points towards the pocket formed by the 23S rRNA residues A2503-U2506. R^-3^ in the SecM(*Ms*) AP remains within this pocket in the MD simulations and appears to form a stable stacking interaction with G2505 (>50% of simulation time in the unperturbed G^-12^ system; Fig. S13).

The mutations in position -12 fall in the proposed regulator module. Our MD results show opposite behaviors of the N^-12^ and K^-12^ systems. On the one hand, we observe stabilizing interactions and a high correlation between N^-12^ and the uL22 loop. Furthermore, a stacking interaction observed between H^-7^ and U2609 (Fig. 6) could also contribute to the observed increase in stalling efficiency of the N^-12^ system. On the other hand, we observe increased flexibility for the K^-12^ system with the lysine side-chain moving between different interaction sites, none of which is particularly stable, in line with the low stalling efficiency seen for the K^-12^ mutant.

Our results demonstrate that FPA is sufficiently sensitive to detect position- and residue-specific differences in how individual amino acids in a nascent chain interact with the ribosomal exit tunnel, and that all-atom MD simulations – despite the limitations on sampling imposed by the large size of the system – may suggest relevant interactions. Further experimental and computational studies including, e.g., additional amino acid residue types and residue combinations, could conceivably further improve our understanding of the physico-chemical characteristics of the exit tunnel environment.

## Data availability

The data underlying this article are available and deposited in Zenodo: 1) Molecular dynamics simulations, filtered to include all the residues within 1.5 nm of the nascent chain, including the nascent chain and the complete tRNA. The initial models with mutated nascent chains and fixed *cis* peptide bonds are also included (doi.org/10.5281/zenodo.13248385); 2) Setup of Molecular dynamic simulation and model (doi.org/10.5281/zenodo.13340587); 3) Molecular dynamics analysis scripts (doi.org/10.5281/zenodo.13259735); 4) Movies of MD simulations starting from 8qoa and NC mutated to match SecM(Ms) sequence (doi.org/10.5281/zenodo.19010605); 5) Molecular Dynamics input files(doi.org/10.5281/zenodo.19210918). 6) EasyQuant files, Excel sheet with the analysis of the .txt files from EasyQuant, and amino acid sequences of all constructs (10.5281/zenodo.18988399).

## Supplementary Data

Supplementary Figures and Tables are available in the SupplementaryMaterial.pdf

## Author contributions

RK performed the Force Profile Analysis experiments. FPA performed Molecular Dynamics simulations. FPA, RK, ML, GvH discussed the results and commented on the manuscript.

## Acknowledgments

We thank Dr. Rickard Hedman (Stockholm University) for programming and maintenance of the EasyQuant software. We thank Dr. Hanlun Jiang (University of California, Berkeley) for his assistance in remodeling the SecM(*Ms*) nascent chain. Fig 1A and B were made using BioRender.

---

**SI Table 1.**
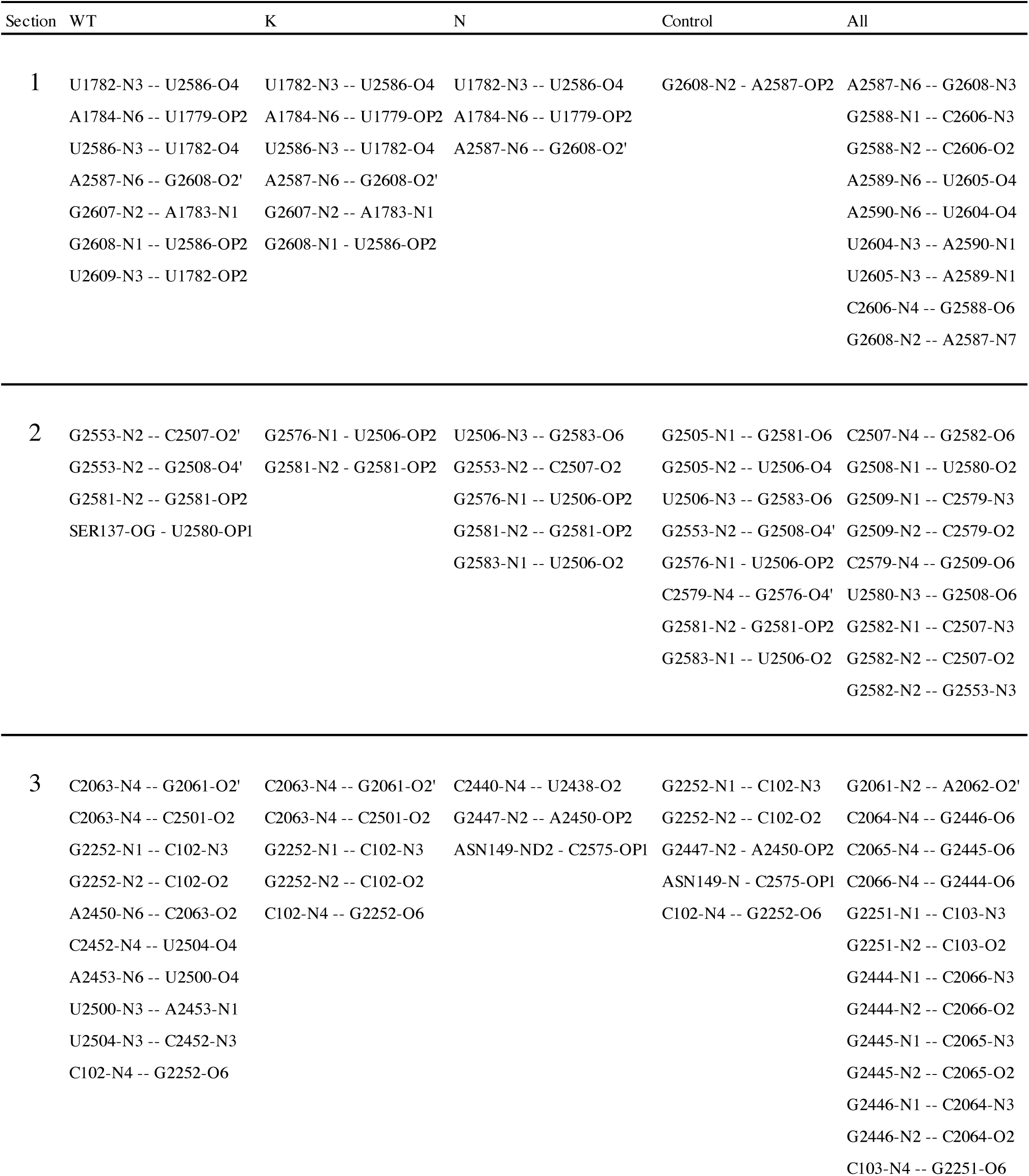
Hydrogen bonds present 50% of the simulation time.

**SI Table 2.**
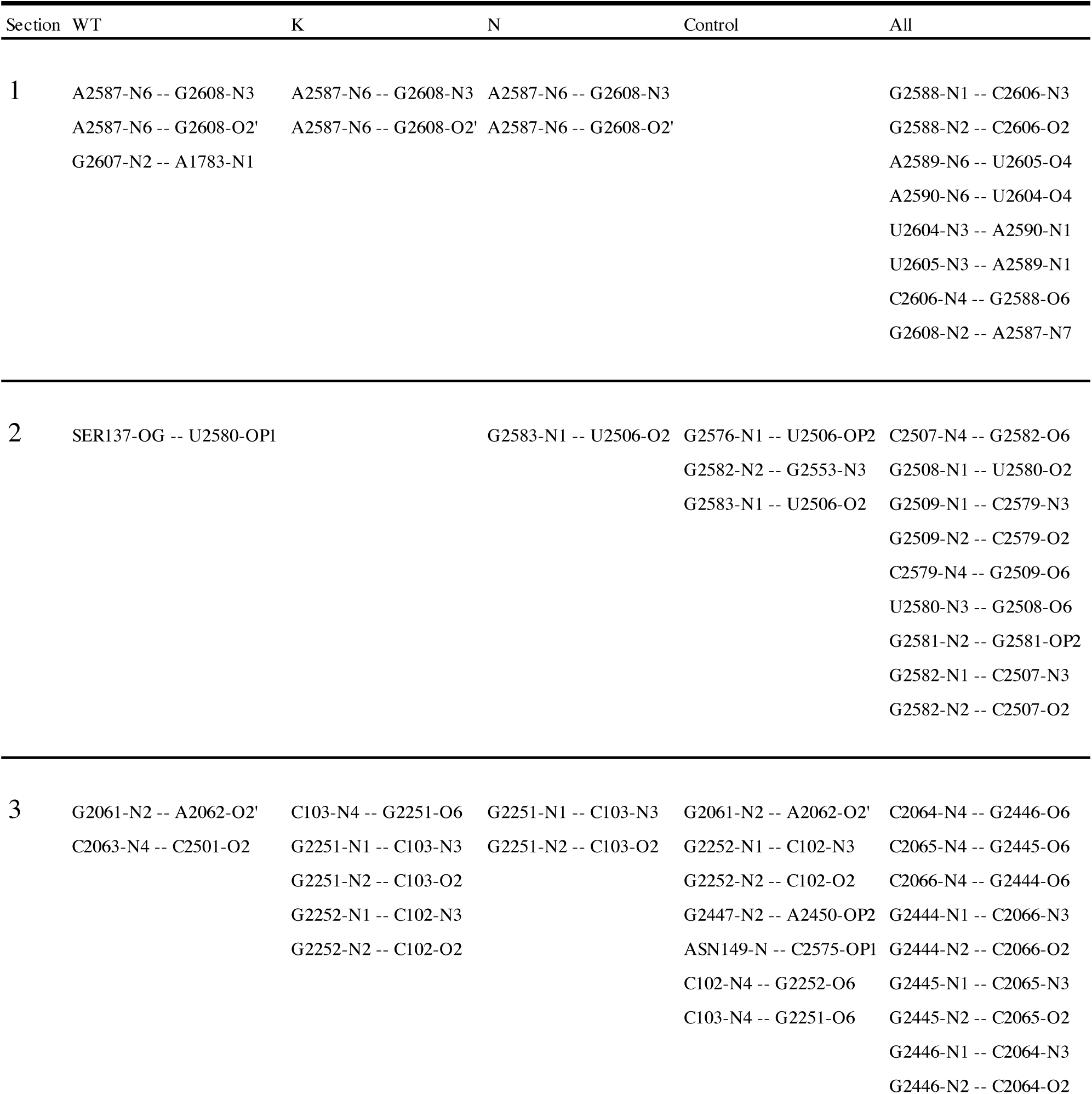
Hydrogen bonds present 75% of the simulation time.

**SI Table 3.**
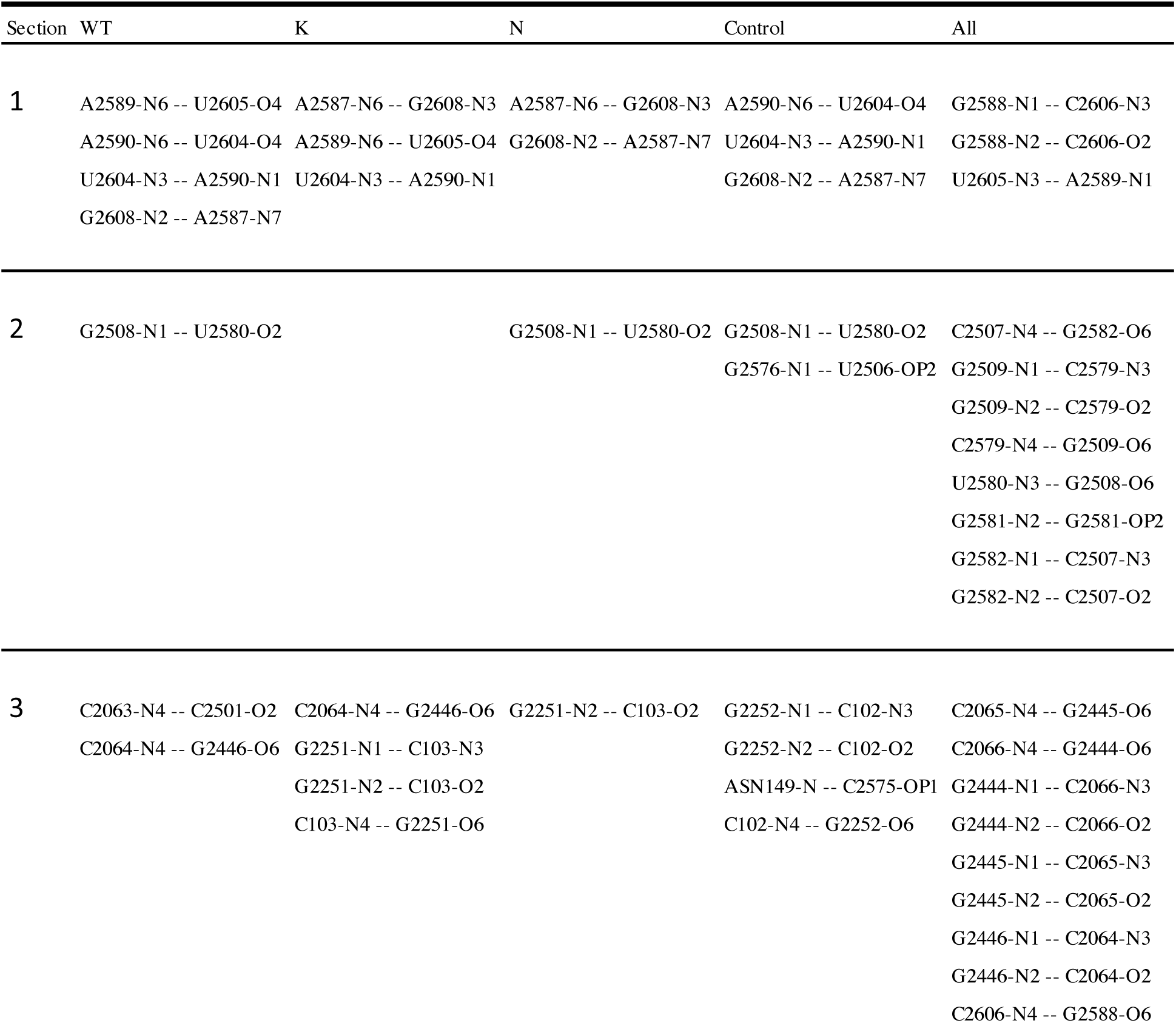
Hydrogen bonds present 90% of the simulation time.

**SI Figure 1.**
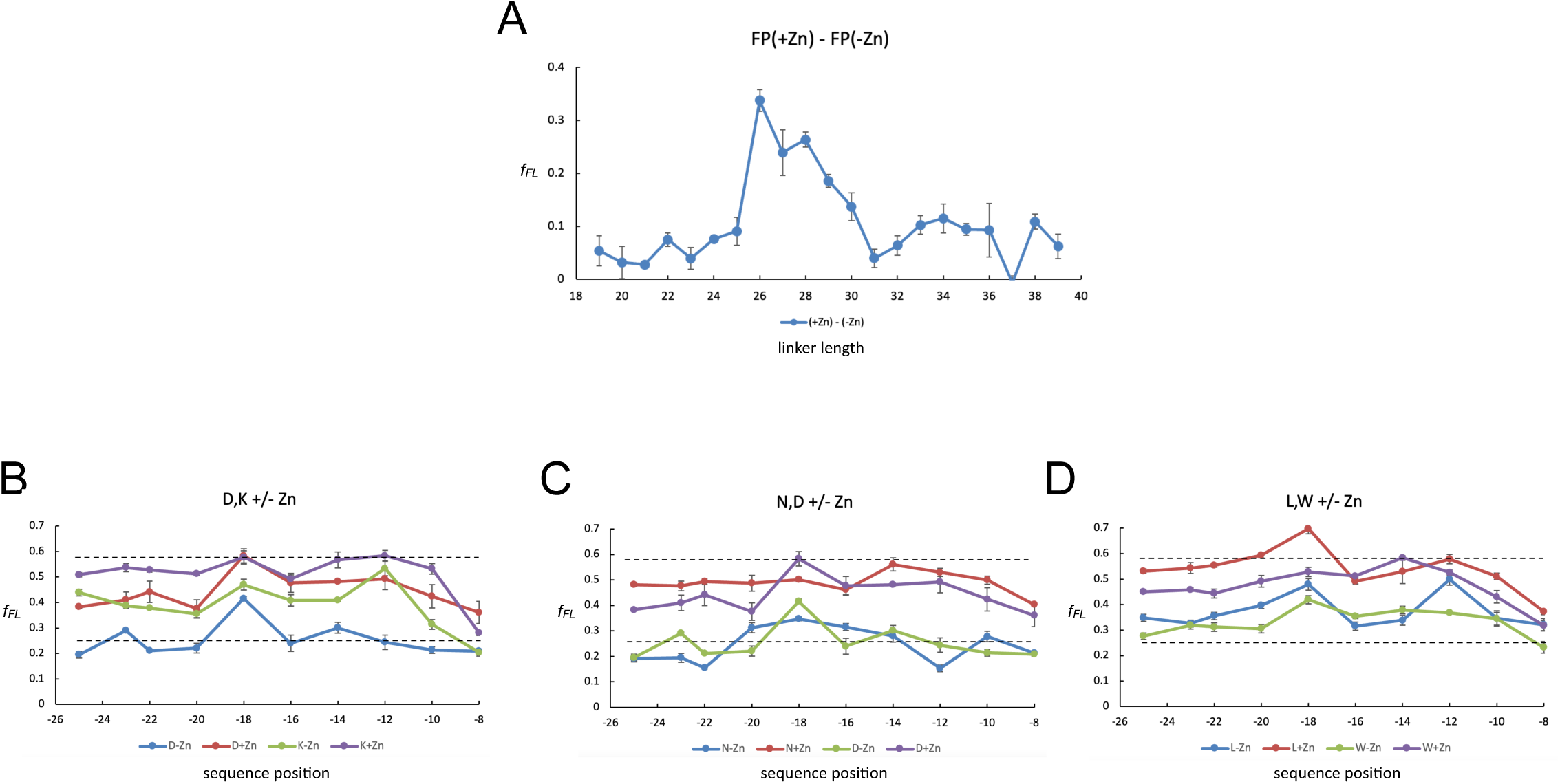
(A) Difference plot of the force profiles obtained ±Zn^2+^ for ADR1a-(GS)-SecM(*Ms*) constructs with GS-segment lengths ranging from 11 to 31 residues (i.e., total linker lengths of 19-40 residues, including the SecM(*Ms*) AP). (B-D) Comparison of force profiles of constructs with D or K (B), N or D (C), L or W (D) at different positions within the GS-segment.

**SI Figure 2.**
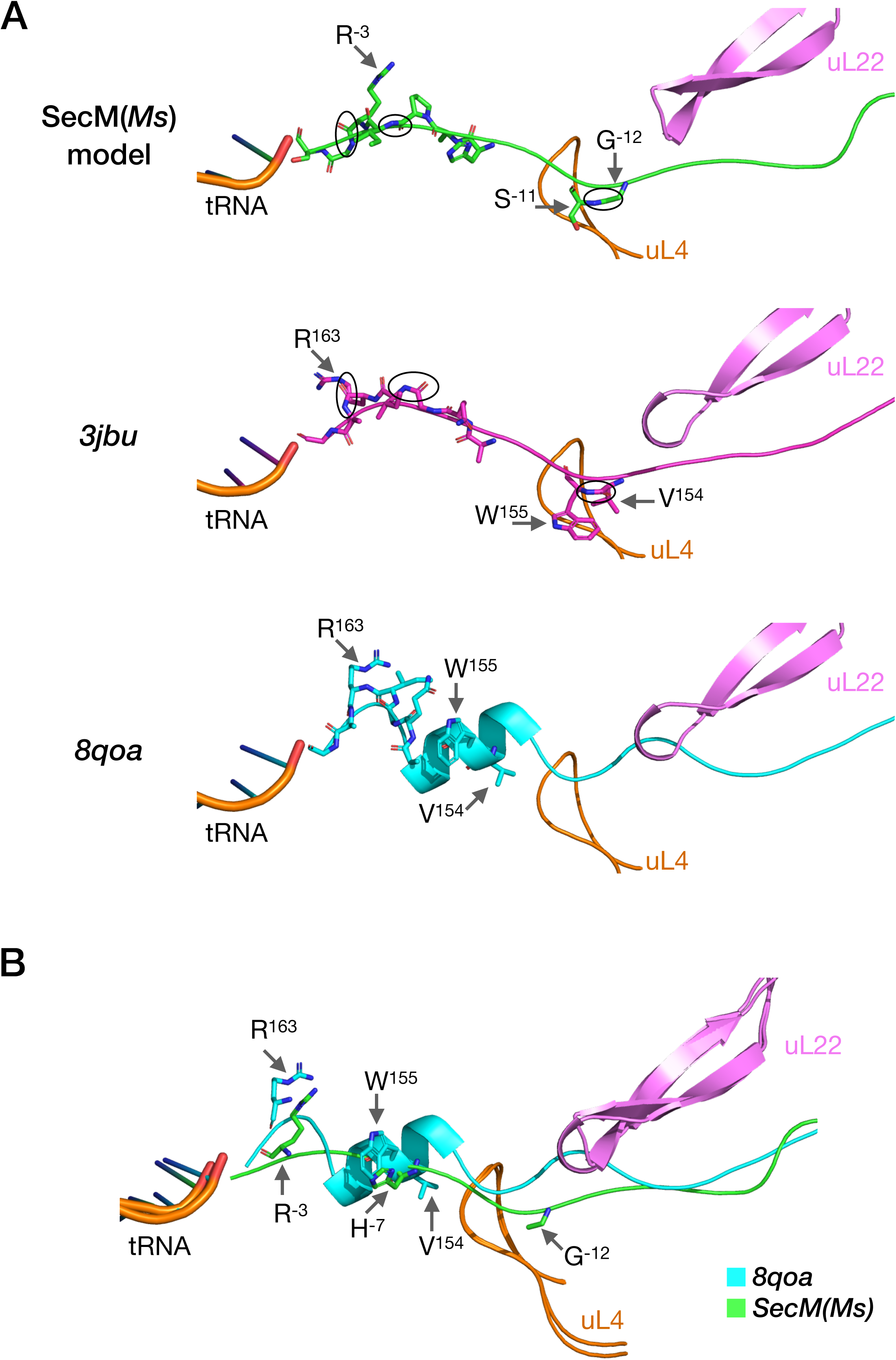
Comparison of the SecM(*Ms*) model used in the MD simulations to SecM(*Ec*) structures *3jbu* and *8qoa*. (A) For each structure, the nascent chain (SecM(*Ms*): green, *3jbu*: magenta, *8qoa*: cyan), tRNA fragment (orange), loops of uL4 (orange) and uL22 (pink) are shown as cartoons. The residues in positions -1 to -7, -11, and -12 (according to the SecM(*Ms*) sequence) are shown as sticks for all structures. For the SecM(*Ms*) model and *3jbu*, the peptide bonds that were remodeled (to correct the *cis* configurations in *3bju*) are circled. Three residues corresponding to the same position in the sequence are labeled: R^-3^/R^163^, S^-11^/W^155^, and G^-12^/V^154^. G^-12^ is the residue mutated to K or N in our simulations. (B) Superposition of the SecM(*Ms*) model (green) and *8qoa* (cyan). Two residues corresponding to the same position in the sequence are shown as sticks and labeled: R^-3^/R^163^, and G^-12^/V^154^. H^-7^ is also shown as sticks to show how it is in the same region as W^155^.

**SI Figure 3.**
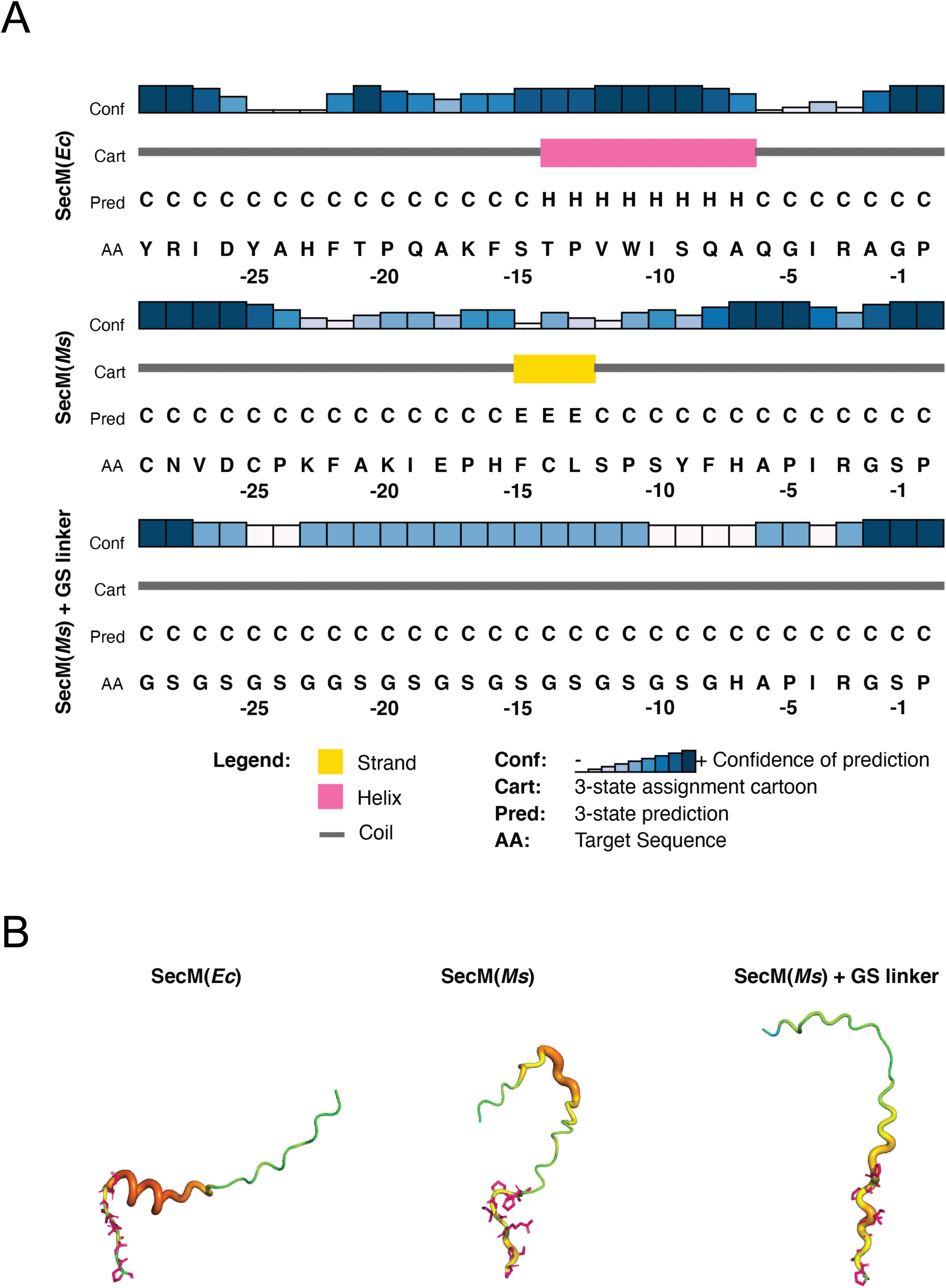
Secondary structure predictions for the AP regions of wildtype SecM(*Ec*), SecM(*Ms*), and the MD model of SecM(*Ms*) + GS segment. (A) PSIPRED prediction (from http://bioinf.cs.ucl.ac.uk/psipred/). The SecM(*Ec*) AP is predicted to contain an α-helix (as observed in the cryo-EM structure), while the SecM(*Ms*) + GS segment is predicted to be a coil. (B) AlphaFold2 predictions from UniProt for SecM(*Ec*) AP (identifier: AF-P62395-F1) and SecM(*Ms*) (identifier:AF-Q65VS7-F1). SecM(*Ms*) + GS segment structure was predicted by ColabFold. Only the amino acids corresponding to the sequence used for the PSIDPRED prediction are shown. The images were generated with PyMol, using the preset visualization “b factor putty”, where instead of using b-factors, the per-residue confidence scores (pLDDT) were used. The structure’s color goes from blue (low confidence) to red (high confidence), and the thickness of the tube also increases with the confidence score. The residues from positions 0 to -7 are shown as pink sticks.

**SI Figure 4.**
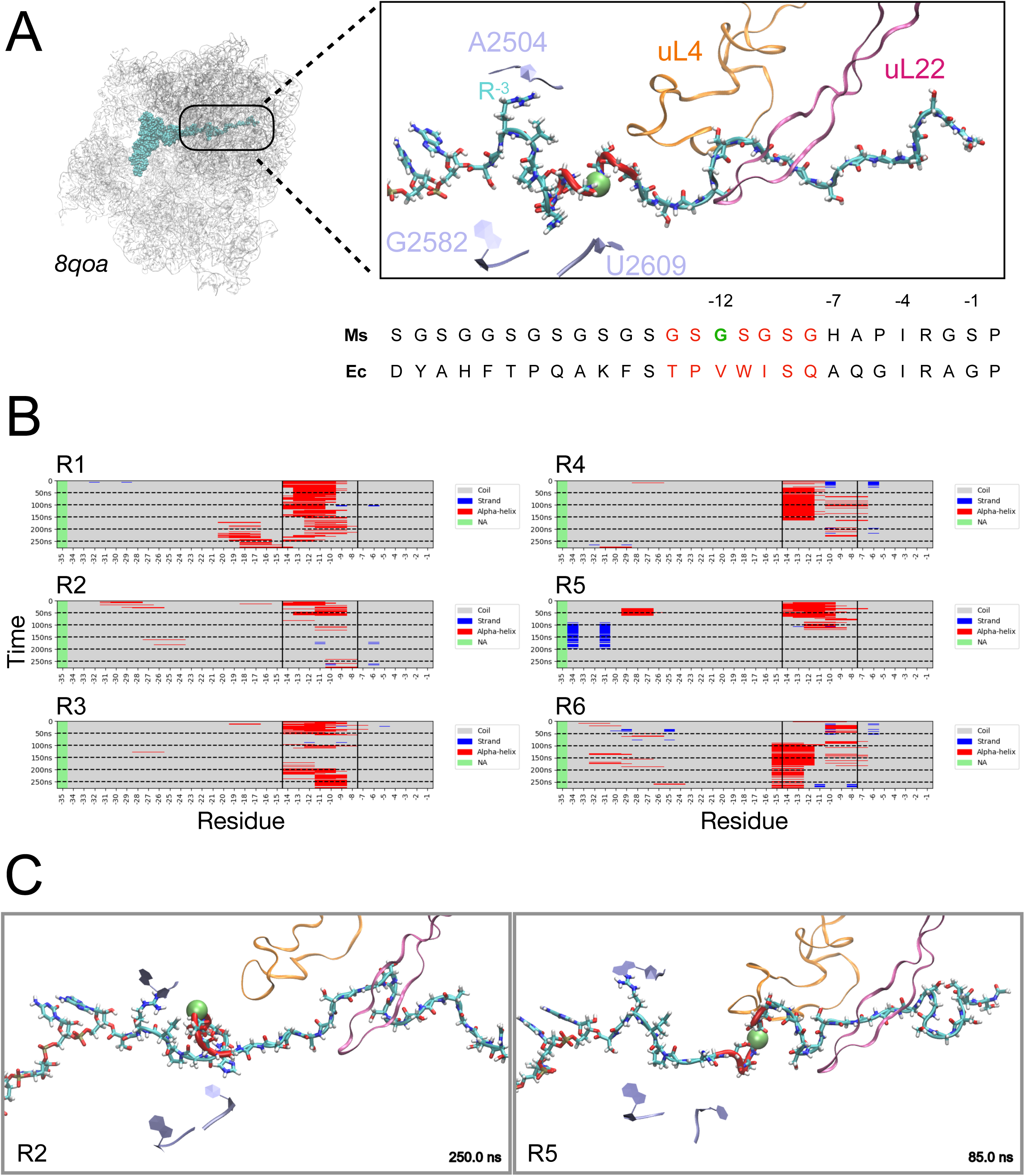
Molecular Dynamics of *8qoa* with the SecM(*Ms*) sequence. (A) Left: The ribosome is shown as a gray cartoon. The tRNA and nascent chain are shown as cyan spheres. The exit tunnel region is highlighted in a square. Right: Close-up of the nascent chain. The sequences of the NC of SecM(*Ec*) and SecM(*Ms*) are shown. The NC and part of the tRNA are shown as a cyan cartoon and licorice representations. The residue at position -12 is shown as a green sphere, and the residues involved in the α-helix observed in *8qoa* are shown in red. The colors of the highlighted residues match between the structure and the sequence. Nucleotides A2501, G2582, and U2609 are shown in light blue for reference to the tunnel orientation. The loops of the uL4 and uL22 proteins are shown in orange and magenta, respectively, for reference. (B) Secondary structure prediction of the NC for the 6 molecular dynamics simulations. The helical region in the SecM(*Ec*) AP in *8qoa* is delimited by vertical black lines. The colors correspond to the secondary structure prediction: coil: gray, strand: blue, ⍺-helix: red, NA: green. (C) Two frames were selected to show the conformations sampled by the NC. The colors match panel A. Movies of the simulations are also provided.

**SI Figure 5.**
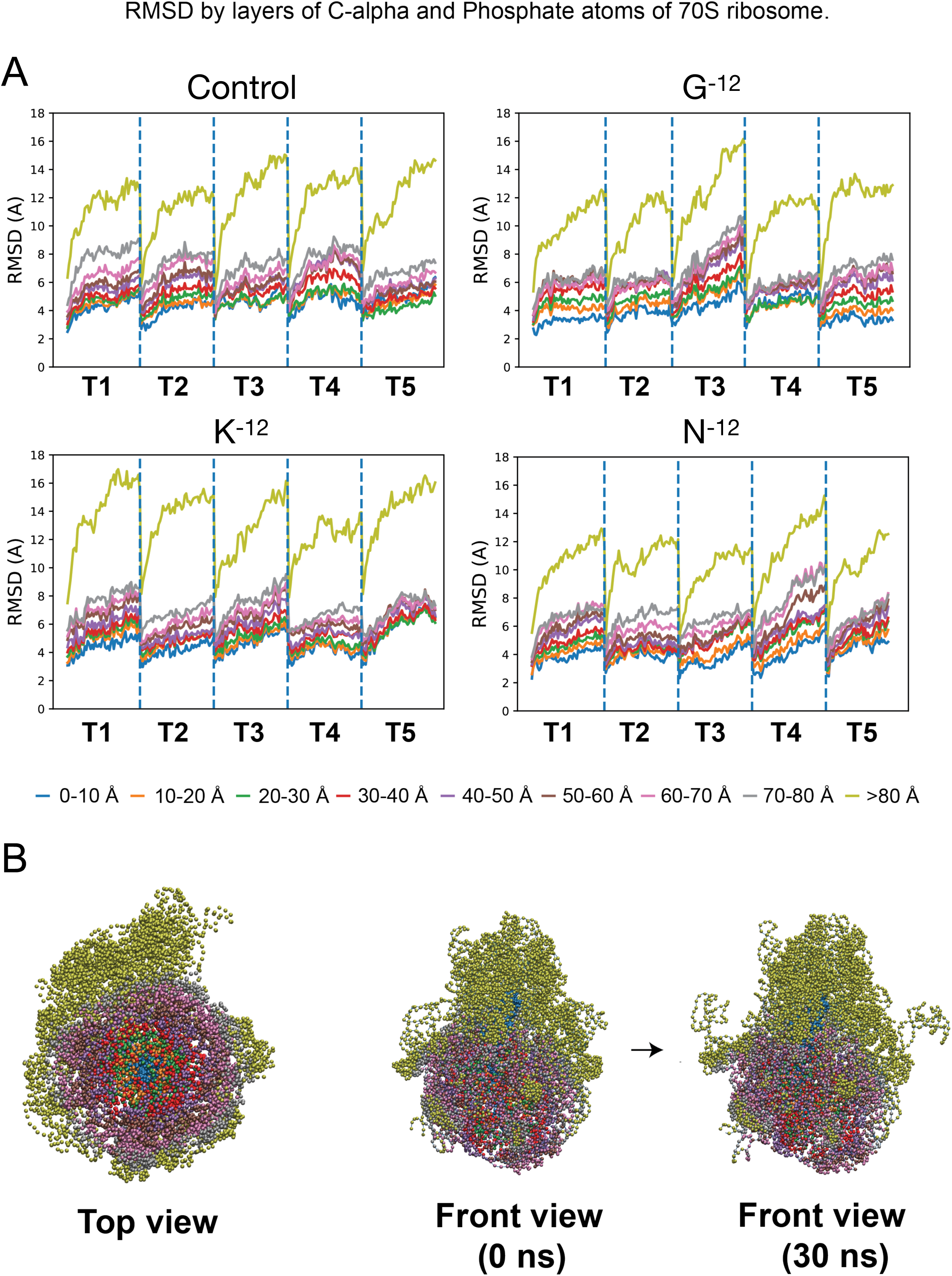
(A) R.M.S.D. of the ribosome by layers. The ribosome was divided into concentric 10 Å layers starting from the nascent peptide and r.m.s.d. values were calculated against each system’s initial model (model after energy minimization). Each plot shows the r.m.s.d traces for each of the five trajectories (T1 to T5) for each system (Control, G^-12^, K^-12^, and N^-12^). (B) Ribosome structure filtered to show only P atoms (for nucleic acids) and C_α_ atoms (for proteins). The top view facilitates the visualization of the concentric layers used to calculate the r.m.s.d in A. The high mobility of the L-1 stalk is reflected in the high r.m.s.d. values observed for the outermost layer (green line), and can be seen in the front-view comparisons between conformations at 0 ns and 30 ns.

**SI Figure 6.**
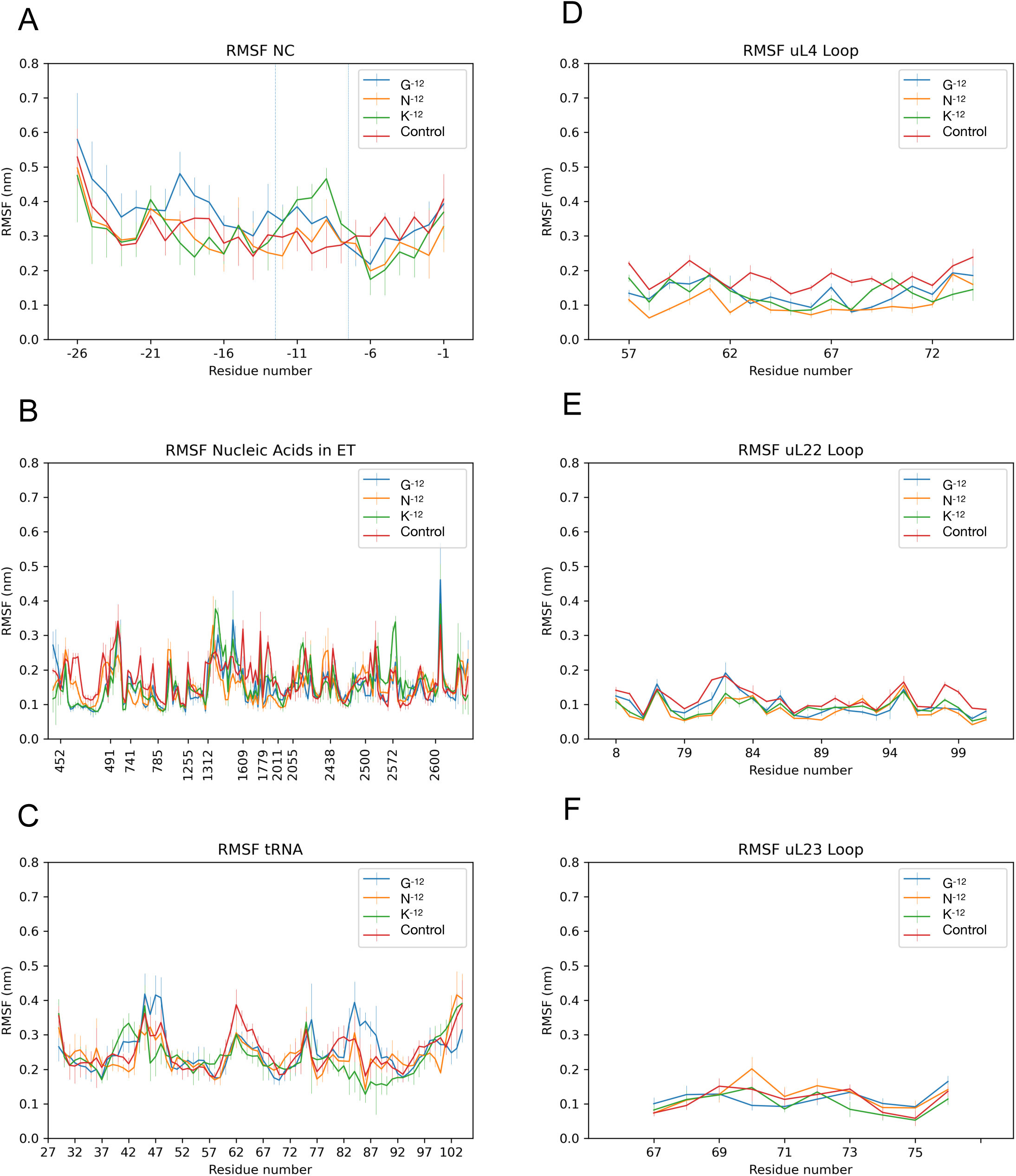
R.M.S.F. of the center of mass of amino acids and nucleic acids in the exit tunnel. Each nucleic acid or amino acid was represented as a bead located in the position of the center of mass (c.o.m.) of each residue. Traces are shown for each of the four systems system. The r.m.s.f values and standard deviation were calculated after bootstrapping. (A) NC Chain. (B) nucleic acids in the ET. (C) tRNA. (D) uL4 loop. (E) uL22 loop. (F) uL23 loop.

**SI Figure 7.**
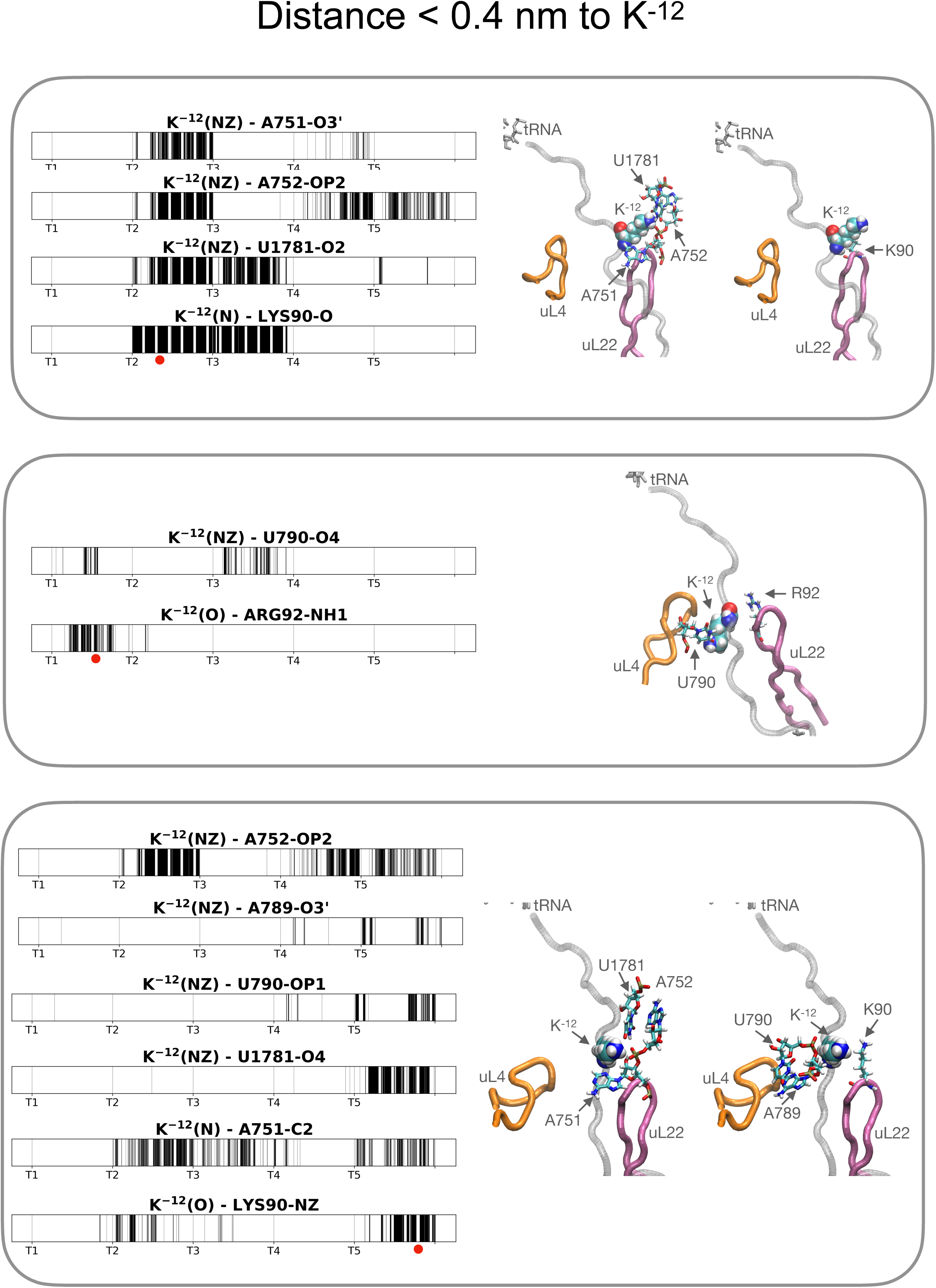
Distance Analysis of the K^-12^ system. Each plot shows a black line when the two atoms specified in the title are within 4 Å. Five trajectories are displayed in each plot, and the beginning of each trajectory is labelled (T1 to T5). The interactions are grouped by similarity to facilitate the structure visualization. Selected frames are shown on the right. The uL4 loop is shown as an orange tube, the uL22 loop is shown as a pink tube, the NC is shown as a grey tube, the tRNA is shown as grey sticks (licorice), and K^-12^ is shown in VdW representation. The interacting residues are shown as sticks (licorice). The red circle under the plot shows the time points from which the frames were extracted. In the top and bottom panels, one frame is shown twice, displaying different interactions to facilitate visualization.

**SI Figure 8.**
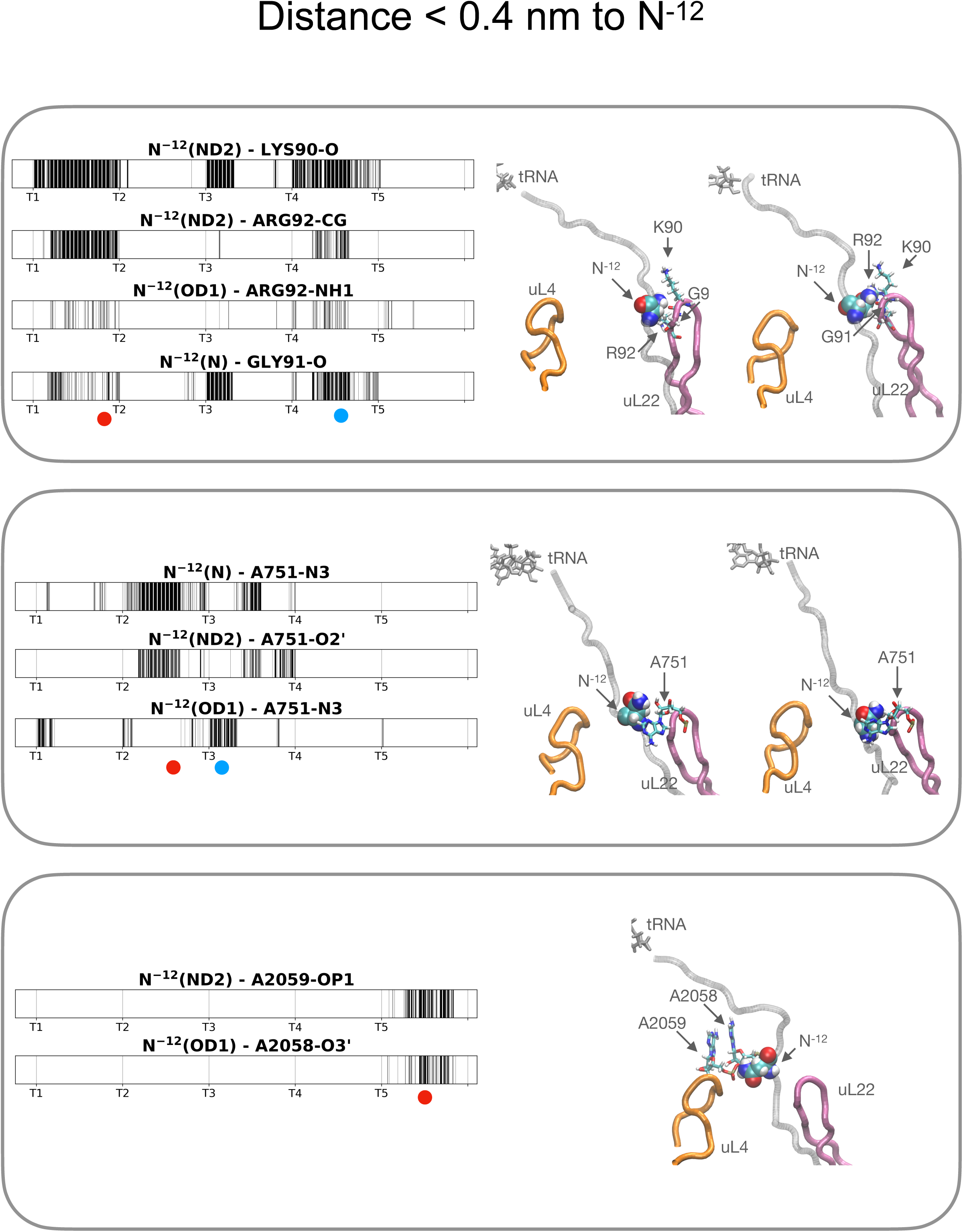
Distance Analysis of N^-12^ system. Each plot shows a black line when the two atoms specified in the title are within 4 Å. Five trajectories are displayed in each plot, and the beginning of each trajectory is labelled (T1 to T5). The interactions are grouped by similarity to facilitate the visualization of the structures. Selected frames are shown on the right. The uL4 loop is shown as an orange tube, the uL22 loop is shown as a pink tube, the NC is shown as a grey tube, the tRNA is shown as grey sticks (licorice), and N^-12^ is shown in VdW representation. The interacting residues are shown as sticks (licorice). The red and blue circles under the plot show the time points from which the frames were extracted. The red circle corresponds to the frame on the left, and the blue circle corresponds to the one on the right.

**SI Figure 9.**
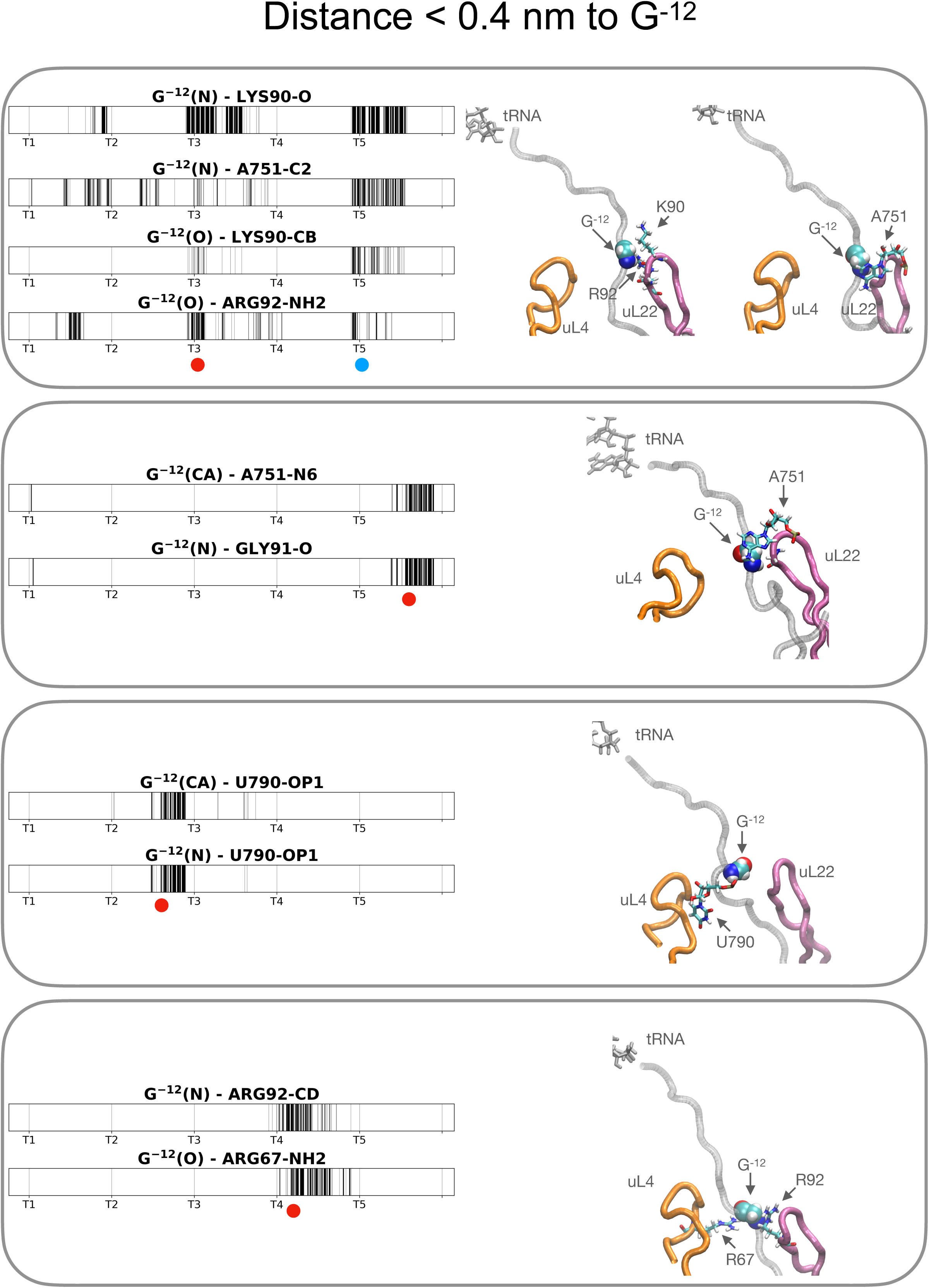
Distance Analysis of G^-12^ (unperturbed) system. Each plot shows a black line when the two atoms specified in the title are within 4 Å. Five trajectories are displayed in each plot, and the beginning of each trajectory is labelled (T1 to T5). The interactions are grouped by similarity to facilitate the visualization of the structures. Selected frames are shown on the right. The uL4 loop is shown as an orange tube, the uL22 loop is shown as a pink tube, the NC is shown as a grey tube, the tRNA is shown as grey sticks (licorice), and G^-12^ is shown in VdW representation. The interacting residues are shown as sticks (licorice). The red and blue circles under the plot show the time points from which the frames were extracted. The red circle corresponds to the frame on the left, and the blue circle corresponds to the one on the right.

**SI Figure 10.**
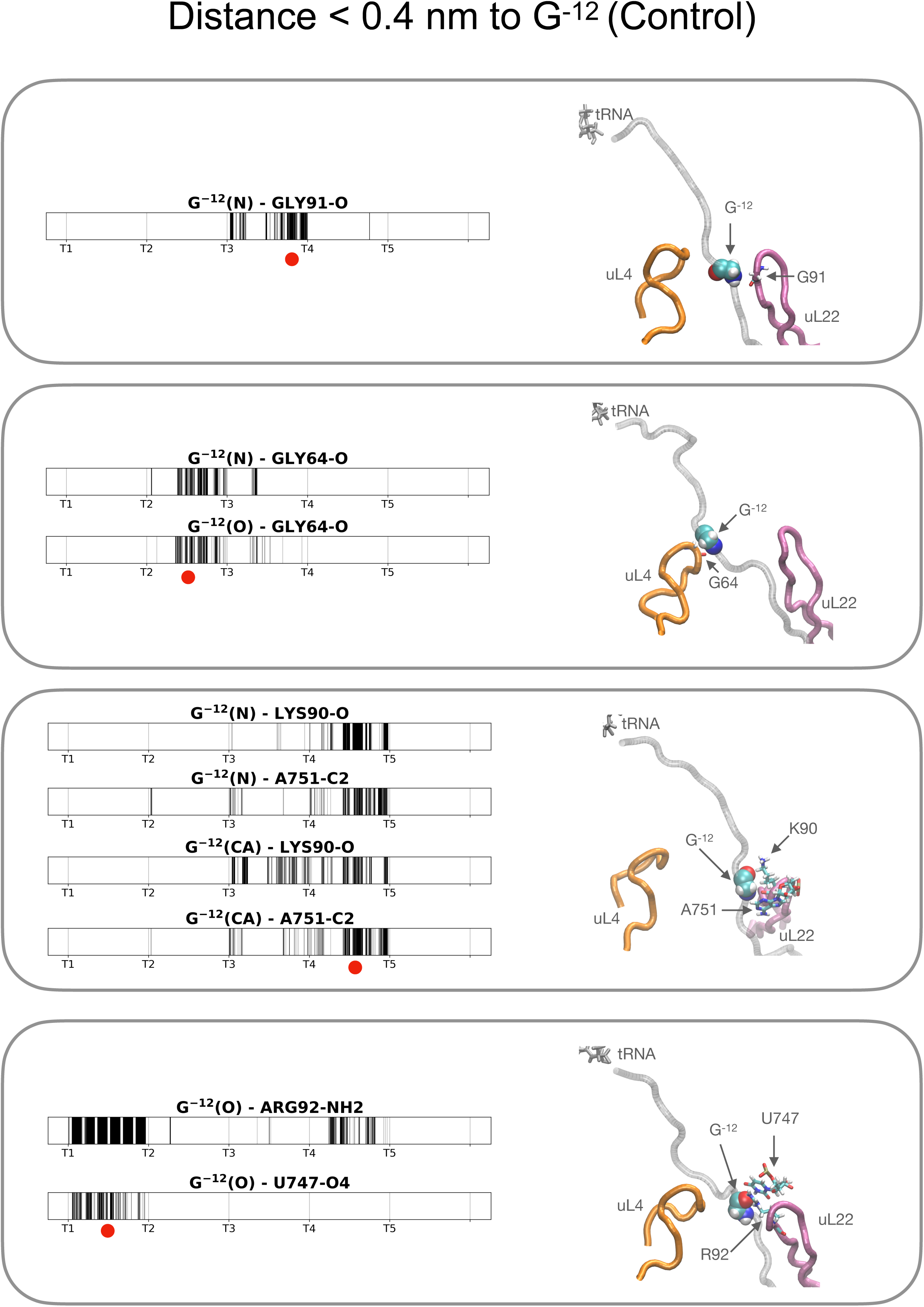
Distance Analysis of G^-12^ (Control) system. Each plot shows a black line when the two atoms specified in the title are within 4 Å. Five trajectories are displayed in each plot, and the beginning of each trajectory is labelled (T1 to T5). The interactions are grouped by similarity to facilitate the visualization of the structures. Selected frames are shown on the right. The uL4 loop is shown as an orange tube, the uL22 loop is shown as a pink tube, the NC is shown as a grey tube, the tRNA is shown as grey sticks (licorice), and G^-12^ is shown in VdW representation. The interacting residues are shown as sticks (licorice). The red and blue circles under the plot show the time points from which the frames were extracted. The red circle corresponds to the frame on the left, and the blue circle corresponds to the one on the right.

**SI Figure 11.**
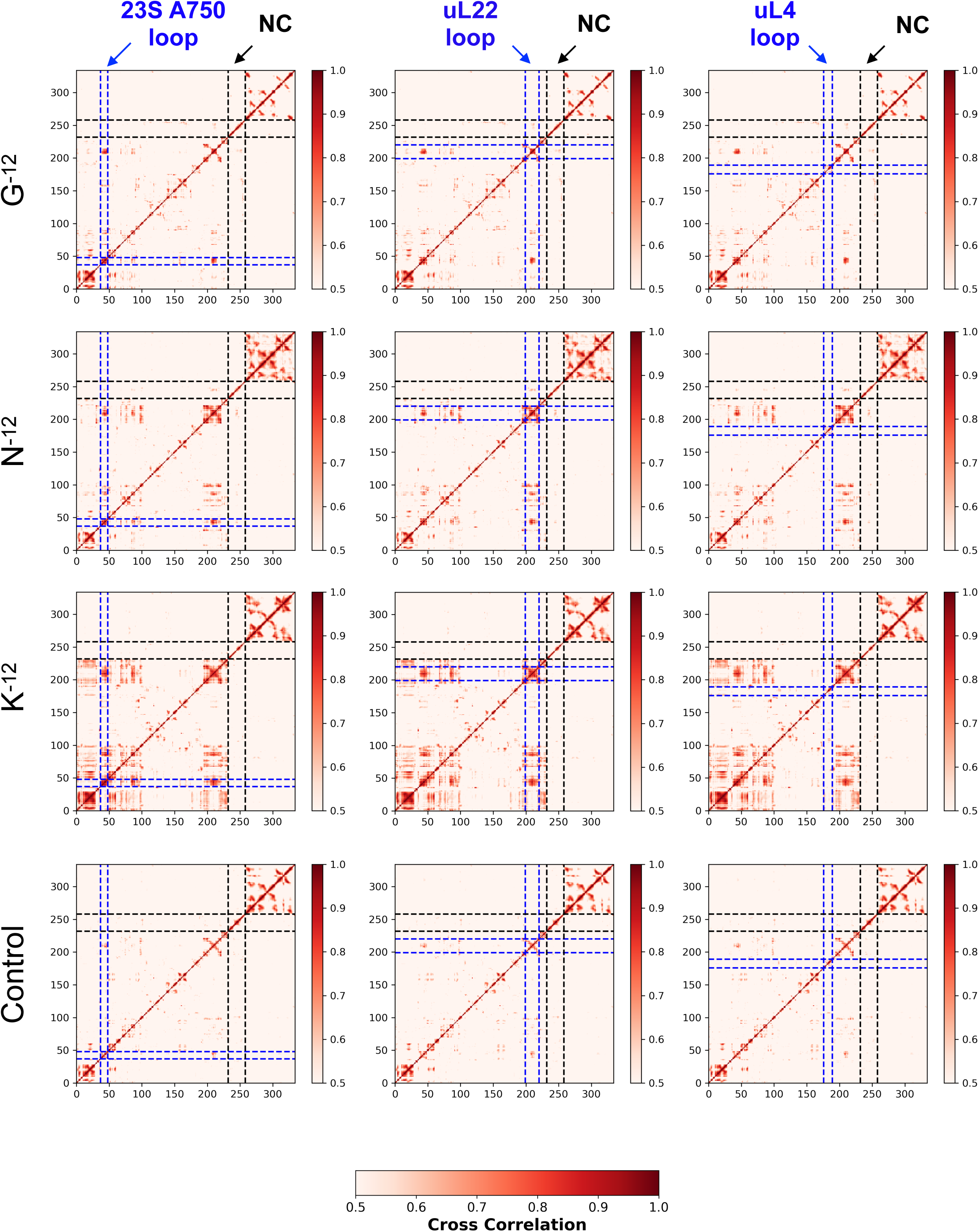
Cross-correlation (Pearson correlation of the covariance matrix) matrices of the nascent chain (NC_, tRNA, and residues within 15 Å of the NC (in total 334 residues). To facilitate the visualization of highly correlated regions, only values higher than 0.5 are shown (refer to color bar). Each row shows the cross-correlation matrix for each of the four simulated systems (WT(G^-12^), N^-12^, K^-12^, and Control). The three columns show the same matrix for each system, but three different regions are highlighted by dashed blue lines: 23S A750 loop (residues 37 to 48 of the small sub-system), uL22 loop (residues 199 to 220 of the small sub-system), and uL4 loop (residues 176 to 189 of the small sub-system). All matrices highlight the region corresponding to the NC (residues 232 to 258 of the small sub-system) with dashed black lines. For N^-12^ and G^-12^ (unperturbed), there is a strong correlation between the NC and residues in the uL22 (as seen in the intersection between the blue and black highlighted regions in the middle panel), which is not seen for K^-12^ and Control. A close-up of this region is shown in SI Fig. S11.

**SI Figure 12.**
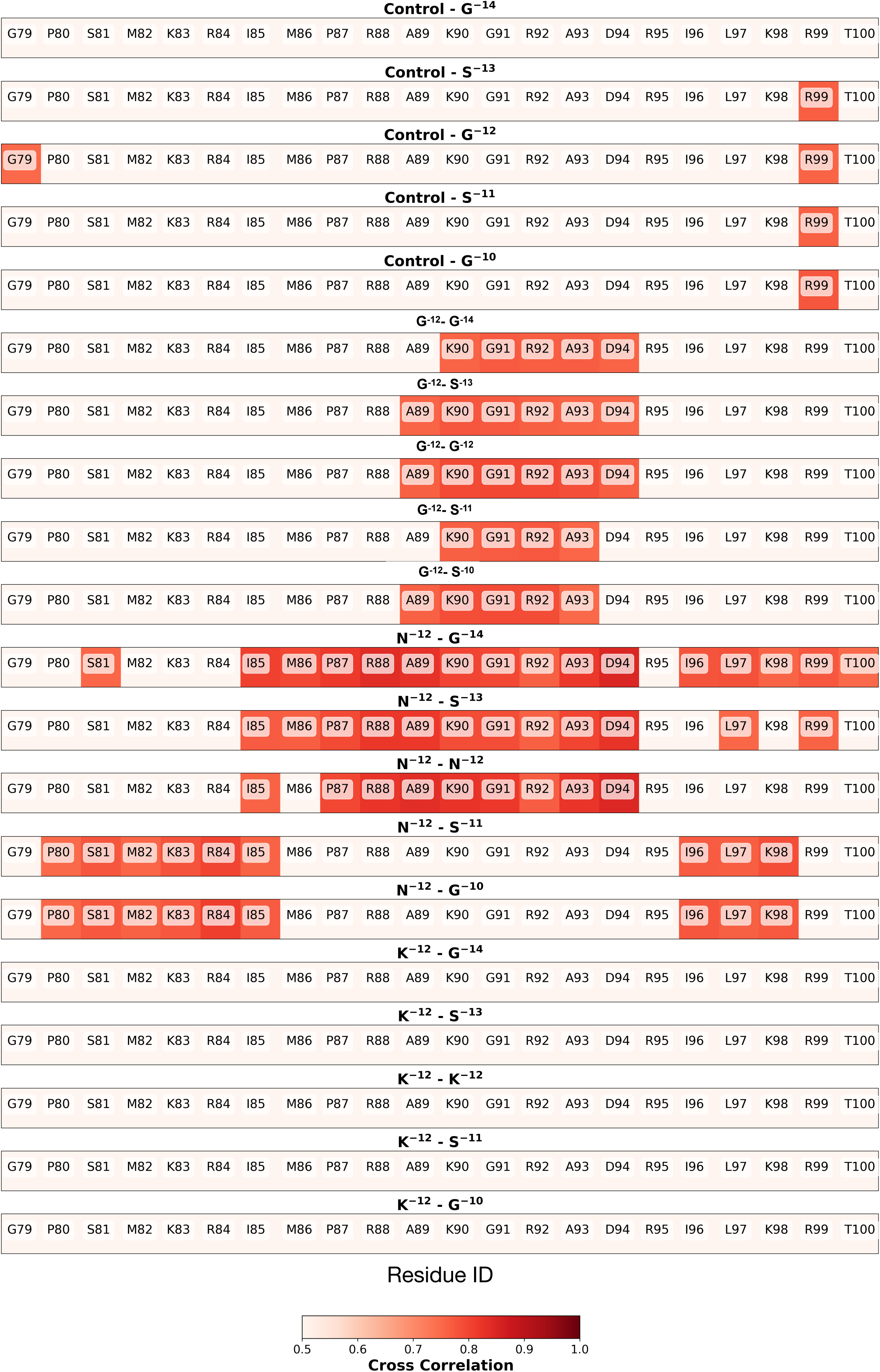
Cross-correlation between the uL22 loop (residues G79 to T100) and residues - 10 to -14 of the nascent chain (NC) for each of the simulated systems ( unperturbed (G^-12^), N^-12^, K^-12^, and Control). To facilitate the visualization of highly correlated regions, only values higher than 0.5 are shown (refer to color bar). The title of each heatmap shows the simulated system and the NC residue for which heatmap is being plotted. Only the unperturbed G^-12^ and N^-12^ systems showed correlation values higher than 0.5 for the tip of the uL22 loop (residues A89 to A93).

**SI Figure 13.**
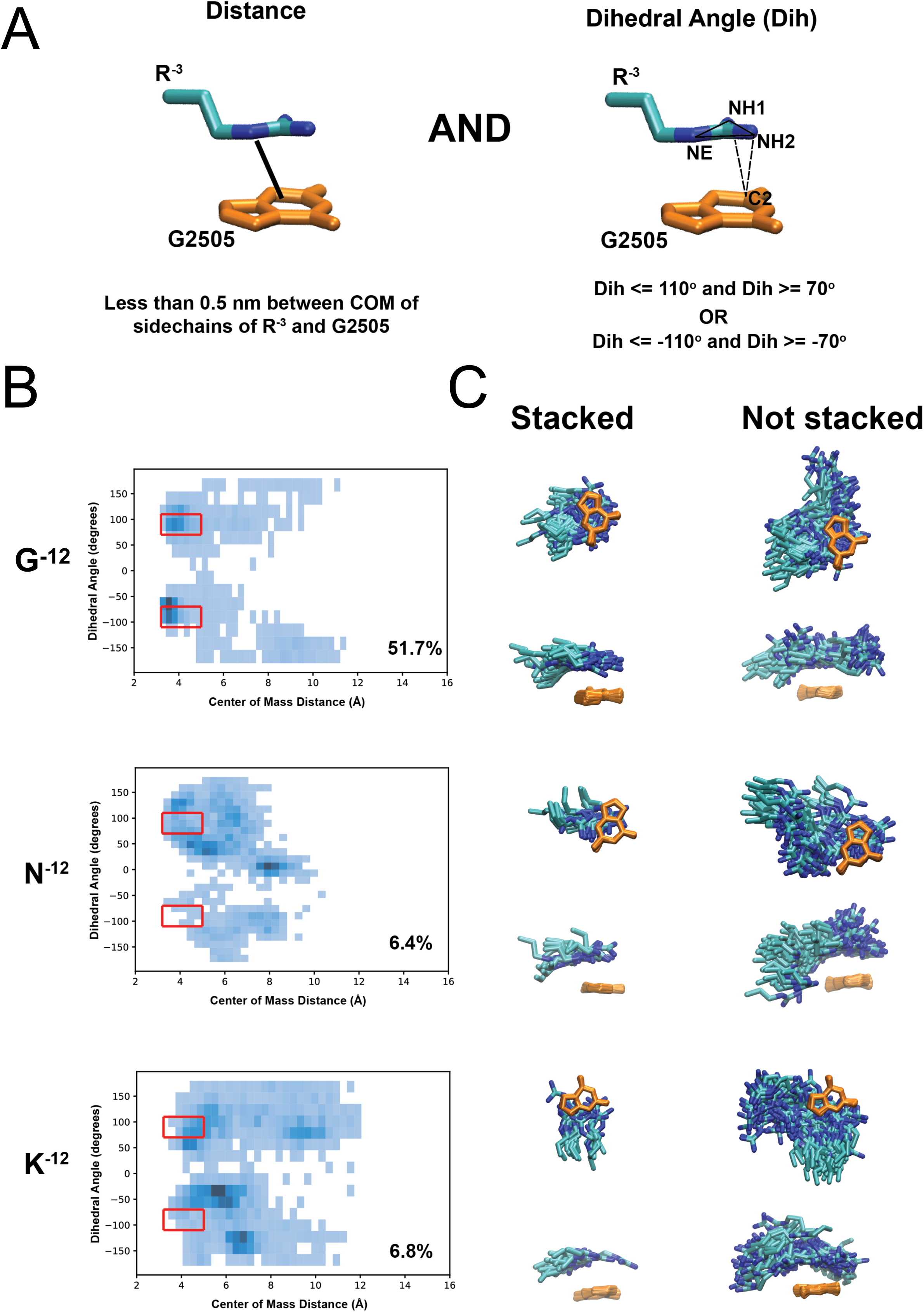
Stacking between R^-3^ and G2505. (A) The two reaction coordinates used to define stacking between R^-3^ and G2505 are shown in the top panel. A stacking interaction is scored if the distance between the center of mass (c.o.m) of the R^-3^ sidechain and the base ring of G2505 is less than 0.5 nm, and the dihedral angle formed between the base ring of G2505 and the C2 atom of R^-3^ is between 70 to 110 degrees. (B) 2D histograms of the projections onto the reaction coordinates described in A. The red rectangles show the regions where the stacking conditions are met, and the percentage of frames that fall within this region is shown in the bottom right of each plot. (C) Extracted structures from the stacked and non-stacked regions highlighted in B. The sidechain of R^-3^ and the base of G2505 are shown as blue and orange sticks, respectively.

