## Supplemental table and figures for "Single-residue effects on the behavior of a nascent polypeptide chain inside the ribosome exit tunnel"

SI Table 1. Hydrogen bonds present 50% of the simulation time

| Section | WT | K | N | Control | All |
|---|---|---|---|---|---|
| 1 | U1782-N3 -- U2586-O4 | U1782-N3 -- U2586-O4 | U1782-N3 -- U2586-O4 | G2608-N2 - A2587-OP2 | A2587-N6 -- G2608-N3 |
| | A1784-N6 -- U1779-OP2 | A1784-N6 -- U1779-OP2 | A1784-N6 -- U1779-OP2 | | G2588-N1 -- C2606-N3 |
| | U2586-N3 -- U1782-O4 | U2586-N3 -- U1782-O4 | A2587-N6 -- G2608-O2' | | G2588-N2 -- C2606-O2 |
| | A2587-N6 -- G2608-O2' | A2587-N6 -- G2608-O2' | | | A2589-N6 -- U2605-O4 |
| | G2607-N2 -- A1783-N1 | G2607-N2 -- A1783-N1 | | | A2590-N6 -- U2604-O4 |
| | G2608-N1 -- U2586-OP2 | G2608-N1 - U2586-OP2 | | | U2604-N3 -- A2590-N1 |
| | U2609-N3 -- U1782-OP2 | | | | U2605-N3 -- A2589-N1 |
| | | | | | C2606-N4 -- G2588-O6 |
| | | | | | G2608-N2 -- A2587-N7 |
| 2 | G2553-N2 -- C2507-O2' | G2576-N1 - U2506-OP2 | U2506-N3 -- G2583-O6 | G2505-N1 -- G2581-O6 | C2507-N4 -- G2582-O6 |
| | G2553-N2 -- G2508-O4' | G2581-N2 - G2581-OP2 | G2553-N2 -- C2507-O2 | G2505-N2 -- U2506-O4 | G2508-N1 -- U2580-O2 |
| | G2581-N2 -- G2581-OP2 | | G2576-N1 -- U2506-OP2 | U2506-N3 -- G2583-O6 | G2509-N1 -- C2579-N3 |
| | SER137-OG - U2580-OP1 | | G2581-N2 -- G2581-OP2 | G2553-N2 -- G2508-O4' | G2509-N2 -- C2579-O2 |
| | | | G2583-N1 -- U2506-O2 | G2576-N1 - U2506-OP2 | C2579-N4 -- G2509-O6 |
| | | | | C2579-N4 -- G2576-O4' | U2580-N3 -- G2508-O6 |
| | | | | G2581-N2 - G2581-OP2 | G2582-N1 -- C2507-N3 |
| | | | | G2583-N1 -- U2506-O2 | G2582-N2 -- C2507-O2 |
| | | | | | G2582-N2 -- G2553-N3 |
| 3 | C2063-N4 -- G2061-O2' | C2063-N4 -- G2061-O2' | C2440-N4 -- U2438-O2 | G2252-N1 -- C102-N3 | G2061-N2 -- A2062-O2' |
| | C2063-N4 -- C2501-O2 | C2063-N4 -- C2501-O2 | G2447-N2 -- A2450-OP2 | G2252-N2 -- C102-O2 | C2064-N4 -- G2446-O6 |
| | G2252-N1 -- C102-N3 | G2252-N1 -- C102-N3 | ASN149-ND2 - C2575-OP1 | G2447-N2 - A2450-OP2 | C2065-N4 -- G2445-O6 |
| | G2252-N2 -- C102-O2 | G2252-N2 -- C102-O2 | | ASN149-N - C2575-OP1 | C2066-N4 -- G2444-O6 |
| | A2450-N6 -- C2063-O2 | C102-N4 -- G2252-O6 | | C102-N4 -- G2252-O6 | G2251-N1 -- C103-N3 |
| | C2452-N4 -- U2504-O4 | | | | G2251-N2 -- C103-O2 |
| | A2453-N6 -- U2500-O4 | | | | G2444-N1 -- C2066-N3 |
| | U2500-N3 -- A2453-N1 | | | | G2444-N2 -- C2066-O2 |
| | U2504-N3 -- C2452-N3 | | | | G2445-N1 -- C2065-N3 |
| | C102-N4 -- G2252-O6 | | | | G2445-N2 -- C2065-O2 |
| | | | | | G2446-N1 -- C2064-N3 |
| | | | | | G2446-N2 -- C2064-O2 |
| | | | | | C103-N4 -- G2251-O6 |

SI Table 2. Hydrogen bonds present 75% of the simulation time

| Section | WT | K | N | Control | All |
|---|---|---|---|---|---|
| 1 | A2587-N6 -- G2608-N3 | A2587-N6 -- G2608-N3 | A2587-N6 -- G2608-N3 | | G2588-N1 -- C2606-N3 |
| | A2587-N6 -- G2608-O2' | A2587-N6 -- G2608-O2' | A2587-N6 -- G2608-O2' | | G2588-N2 -- C2606-O2 |
| | G2607-N2 -- A1783-N1 | | | | A2589-N6 -- U2605-O4 |
| | | | | | A2590-N6 -- U2604-O4 |
| | | | | | U2604-N3 -- A2590-N1 |
| | | | | | U2605-N3 -- A2589-N1 |
| | | | | | C2606-N4 -- G2588-O6 |
| | | | | | G2608-N2 -- A2587-N7 |
| 2 | SER137-OG -- U2580-OP1 | | G2583-N1 -- U2506-O2 | G2576-N1 -- U2506-OP2 | C2507-N4 -- G2582-O6 |
| | | | | G2582-N2 -- G2553-N3 | G2508-N1 -- U2580-O2 |
| | | | | G2583-N1 -- U2506-O2 | G2509-N1 -- C2579-N3 |
| | | | | | G2509-N2 -- C2579-O2 |
| | | | | | C2579-N4 -- G2509-O6 |
| | | | | | U2580-N3 -- G2508-O6 |
| | | | | | G2581-N2 -- G2581-OP2 |
| | | | | | G2582-N1 -- C2507-N3 |
| | | | | | G2582-N2 -- C2507-O2 |
| 3 | G2061-N2 -- A2062-O2' | C103-N4 -- G2251-O6 | G2251-N1 -- C103-N3 | G2061-N2 -- A2062-O2' | C2064-N4 -- G2446-O6 |
| | C2063-N4 -- C2501-O2 | G2251-N1 -- C103-N3 | G2251-N2 -- C103-O2 | G2252-N1 -- C102-N3 | C2065-N4 -- G2445-O6 |
| | | G2251-N2 -- C103-O2 | | G2252-N2 -- C102-O2 | C2066-N4 -- G2444-O6 |
| | | G2252-N1 -- C102-N3 | | G2447-N2 -- A2450-OP2 | G2444-N1 -- C2066-N3 |
| | | G2252-N2 -- C102-O2 | | ASN149-N -- C2575-OP1 | G2444-N2 -- C2066-O2 |
| | | | | C102-N4 -- G2252-O6 | G2445-N1 -- C2065-N3 |
| | | | | C103-N4 -- G2251-O6 | G2445-N2 -- C2065-O2 |
| | | | | | G2446-N1 -- C2064-N3 |
| | | | | | G2446-N2 -- C2064-O2 |

SI Table 3. Hydrogen bonds present 90% of the simulation time

| Section | WT | K | N | Control | All |
|---|---|---|---|---|---|
| 1 | A2589-N6 -- U2605-O4 | A2587-N6 -- G2608-N3 | A2587-N6 -- G2608-N3 | A2590-N6 -- U2604-O4 | G2588-N1 -- C2606-N3 |
| | A2590-N6 -- U2604-O4 | A2589-N6 -- U2605-O4 | G2608-N2 -- A2587-N7 | U2604-N3 -- A2590-N1 | G2588-N2 -- C2606-O2 |
| | U2604-N3 -- A2590-N1 | U2604-N3 -- A2590-N1 | | G2608-N2 -- A2587-N7 | U2605-N3 -- A2589-N1 |
| | G2608-N2 -- A2587-N7 | | | | |
| 2 | G2508-N1 -- U2580-O2 | | G2508-N1 -- U2580-O2 | G2508-N1 -- U2580-O2 | C2507-N4 -- G2582-O6 |
| | | | | G2576-N1 -- U2506-OP2 | G2509-N1 -- C2579-N3 |
| | | | | | G2509-N2 -- C2579-O2 |
| | | | | | C2579-N4 -- G2509-O6 |
| | | | | | U2580-N3 -- G2508-O6 |
| | | | | | G2581-N2 -- G2581-OP2 |
| | | | | | G2582-N1 -- C2507-N3 |
| | | | | | G2582-N2 -- C2507-O2 |
| 3 | C2063-N4 -- C2501-O2 | C2064-N4 -- G2446-O6 | G2251-N2 -- C103-O2 | G2252-N1 -- C102-N3 | C2065-N4 -- G2445-O6 |
| | C2064-N4 -- G2446-O6 | G2251-N1 -- C103-N3 | | G2252-N2 -- C102-O2 | C2066-N4 -- G2444-O6 |
| | | G2251-N2 -- C103-O2 | | ASN149-N -- C2575-OP1 | G2444-N1 -- C2066-N3 |
| | | C103-N4 -- G2251-O6 | | C102-N4 -- G2252-O6 | G2444-N2 -- C2066-O2 |
| | | | | | G2445-N1 -- C2065-N3 |
| | | | | | G2445-N2 -- C2065-O2 |
| | | | | | G2446-N1 -- C2064-N3 |
| | | | | | G2446-N2 -- C2064-O2 |
| | | | | | C2606-N4 -- G2588-O6 |

**SI Figure legends**

**SI Figure 1.** (A) Difference plot of the force profiles obtained $\pm Zn^{2+}$ for ADR1a-(GS)-SecM(*Ms*) constructs with GS-segment lengths ranging from 11 to 31 residues (i.e., total linker lengths of 19-40 residues, including the SecM(*Ms*) AP). (B-D) Comparison of force profiles of constructs with D or K (B), N or D (C), L or W (D) at different positions within the GS-segment.

**SI Figure 13.** Stacking between $R^{-3}$ and G2505. (A) The two reaction coordinates used to define stacking between $R^{-3}$ and G2505 are shown in the top panel. A stacking interaction is scored if the distance between the center of mass (c.o.m) of the $R^{-3}$ sidechain and the base ring of G2505 is less than 0.5 nm, and the dihedral angle formed between the base ring of G2505 and the C2 atom of $R^{-3}$ is between 70 to 110 degrees. (B) 2D histograms of the projections onto the reaction coordinates described in A. The red rectangles show the regions where the stacking conditions are met, and the percentage of frames that fall within this region is shown in the bottom right of each plot. (C) Extracted structures from the stacked and non-stacked regions highlighted in B. The sidechain of $R^{-3}$ and the base of G2505 are shown as blue and orange sticks, respectively.

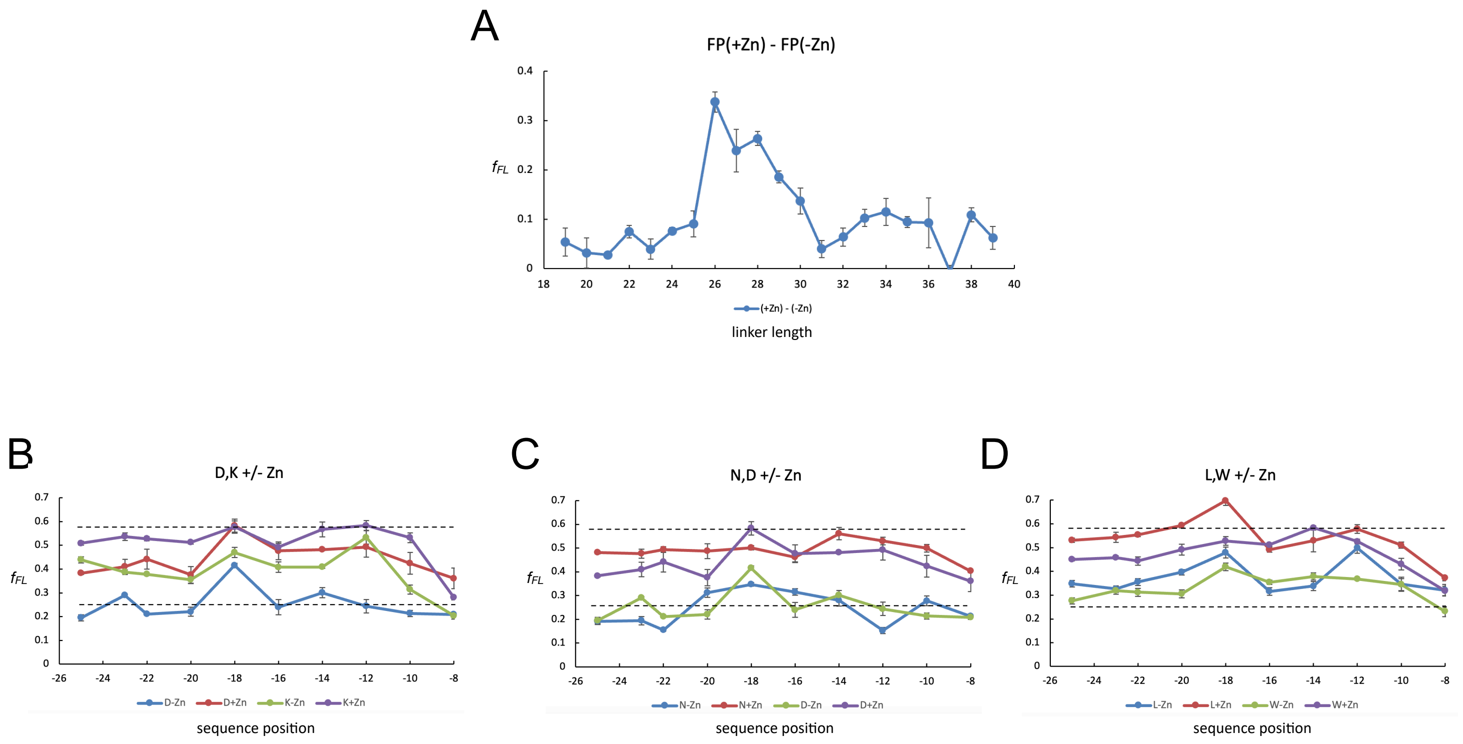
A
FP(+Zn) - FP(-Zn)
fFL
linker length
(+Zn) - (-Zn)
B
D,K +/- Zn
C
N,D +/- Zn
D
L,W +/- Zn
sequence position
D-Zn
D+Zn
K-Zn
K+Zn
N-Zn
N+Zn
D-Zn
D+Zn
L-Zn
L+Zn
W-Zn
W+Zn

Figure S1

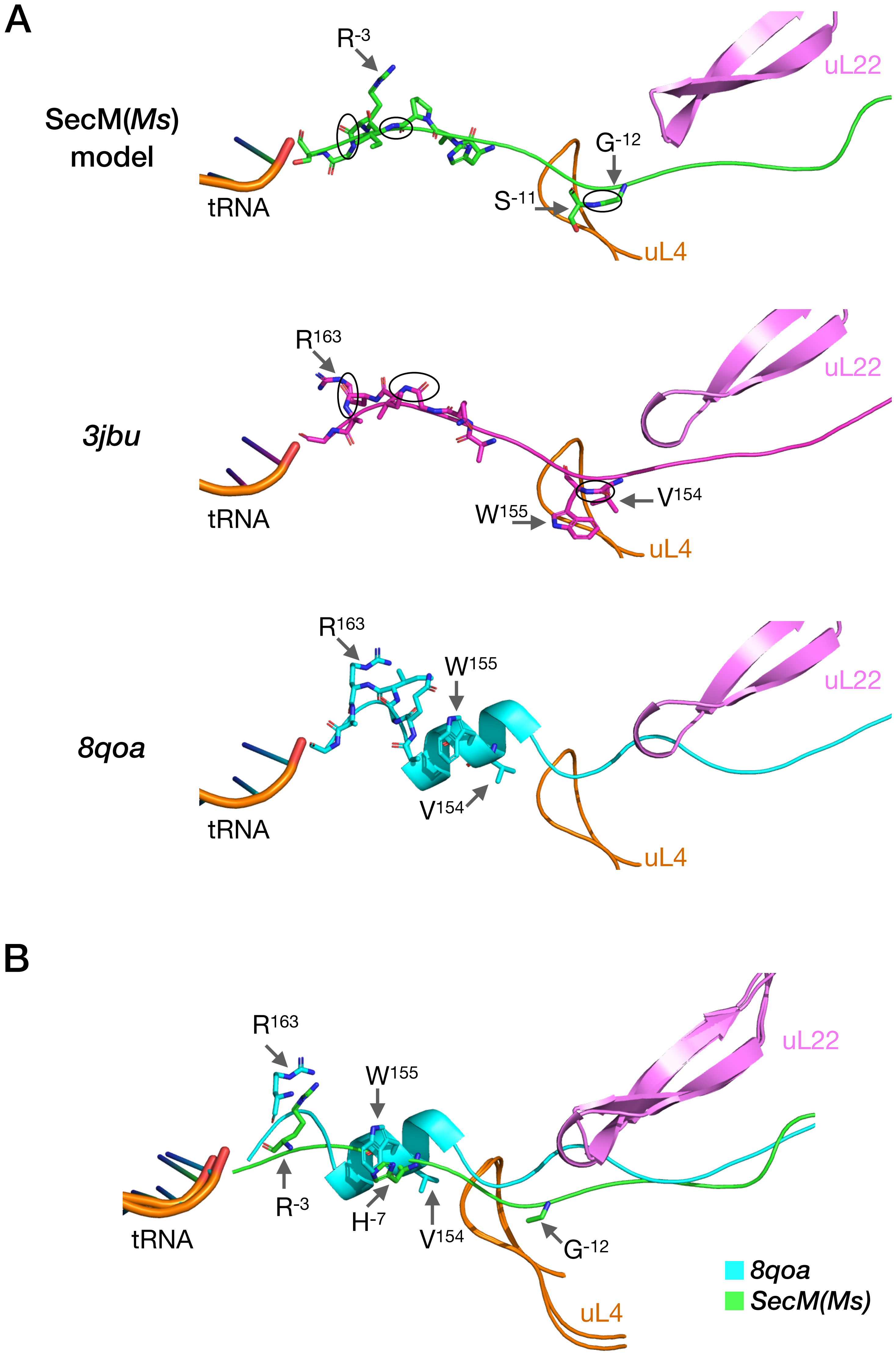

A
R-3
uL22
SecM(Ms)
model
G-12
S-11
tRNA
uL4
R163
uL22
3jbu
V154
W155
tRNA
uL4
R163
W155
uL22
8qoa
V154
tRNA
uL4
B
R163
W155
uL22
R-3
H-7
V154
G-12
tRNA
uL4
8qoa
SecM(Ms)


Figure S2

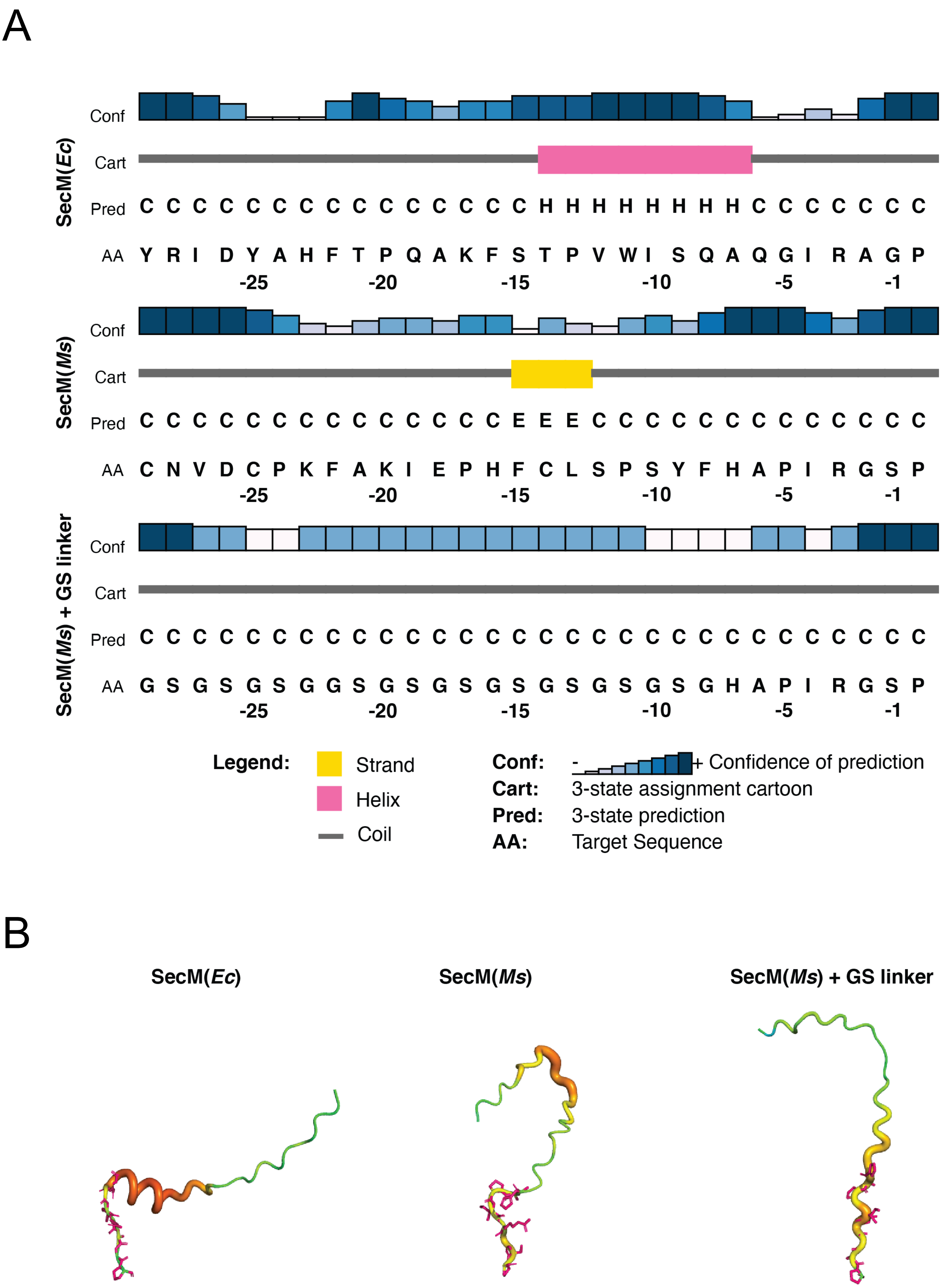
A
SecM(Ec)
Conf
Cart
Pred C C C C C C C C C C C C C C H H H H H H H C C C C C C C
AA Y R I D Y A H F T P Q A K F S T P V W I S Q A Q G I R A G P
-25 -20 -15 -10 -5 -1
SecM(Ms)
Conf
Cart
Pred C C C C C C C C C C C C C C E E E C C C C C C C C C C C C C
AA C N V D C P K F A K I E P H F C L S P S Y F H A P I R G S P
-25 -20 -15 -10 -5 -1
SecM(Ms) + GS linker
Conf
Cart
Pred C C C C C C C C C C C C C C C C C C C C C C C C C C C C C C
AA G S G S G S G G S G S G S G S G S G S G S G H A P I R G S P
-25 -20 -15 -10 -5 -1
Legend:
Strand
Helix
Coil
Conf: - + Confidence of prediction
Cart: 3-state assignment cartoon
Pred: 3-state prediction
AA: Target Sequence
B
SecM(Ec)
SecM(Ms)
SecM(Ms) + GS linker

Figure S3

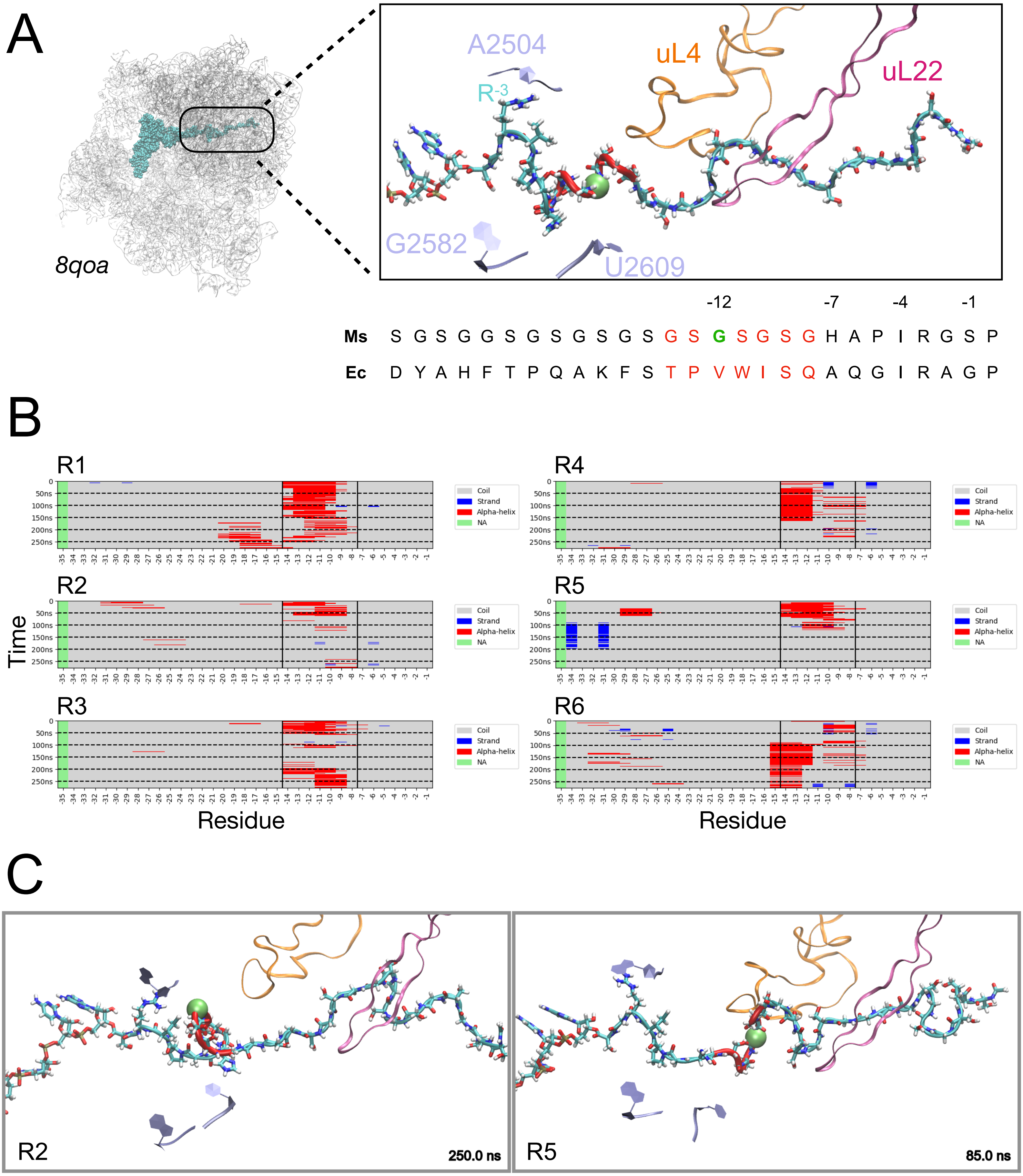
A
8qoa
A2504
R-3
uL4
uL22
G2582
U2609
-12
-7
-4
-1
Ms S G S G G S G S G S G S G S G S G S G H A P I R G S P
Ec D Y A H F T P Q A K F S T P V W I S Q A Q G I R A G P
B
R1
R2
R3
R4
R5
R6
Time
Residue
Coil
Strand
Alpha-helix
NA
C
R2
250.0 ns
R5
85.0 ns

Figure S4

RMSD by layers of C-alpha and Phosphate atoms of 70S ribosome.

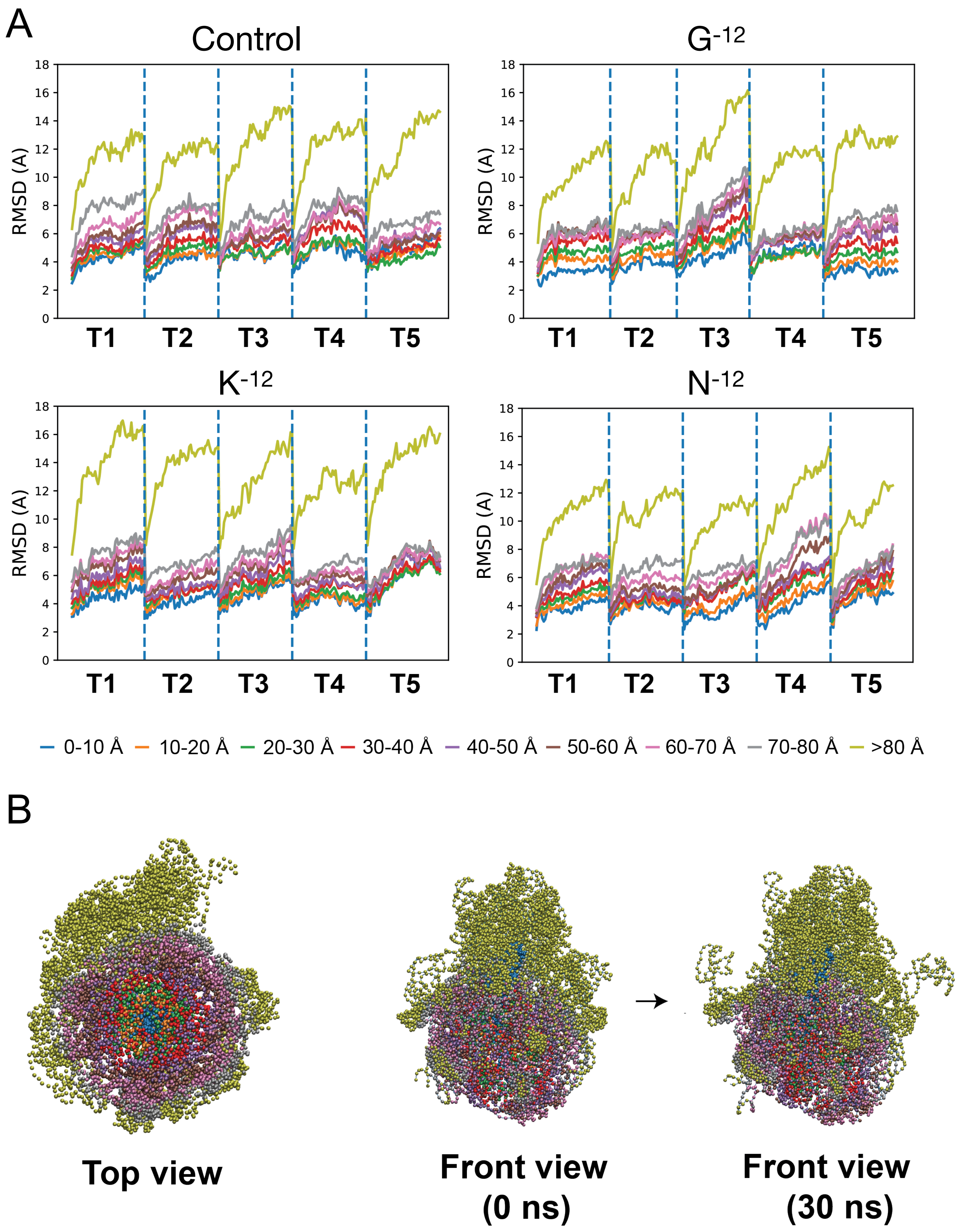


Figure S5

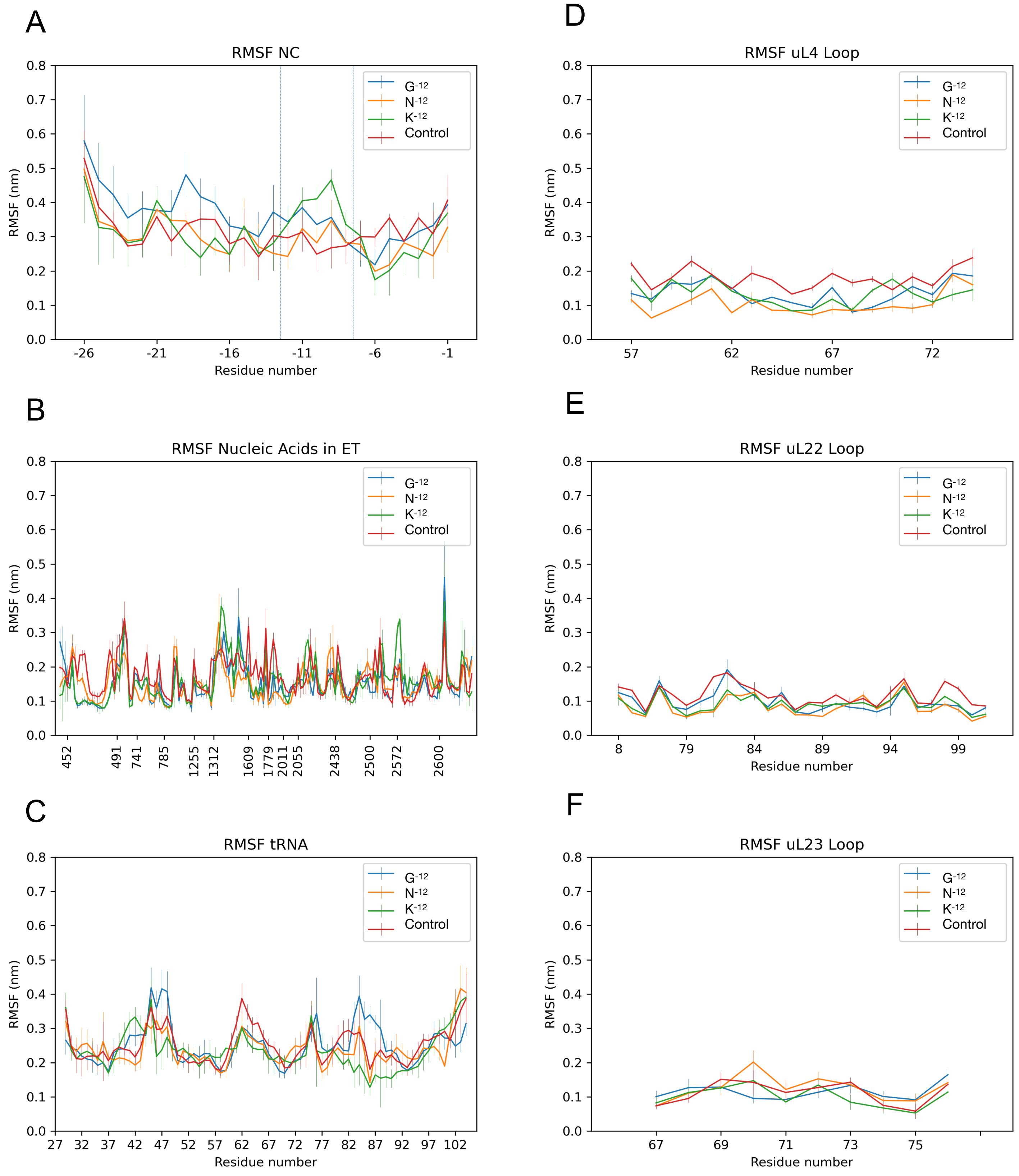
A
RMSF NC
RMSF (nm)
Residue number
G$^{-12}$
N$^{-12}$
K$^{-12}$
Control
B
RMSF Nucleic Acids in ET
RMSF (nm)
452 491 741 785 1255 1312 1609 1779 2011 2055 2438 2500 2572 2600
C
RMSF tRNA
RMSF (nm)
Residue number
D
RMSF uL4 Loop
RMSF (nm)
Residue number
E
RMSF uL22 Loop
RMSF (nm)
Residue number
F
RMSF uL23 Loop
RMSF (nm)
Residue number

Figure S6

### Distance < 0.4 nm to K-12

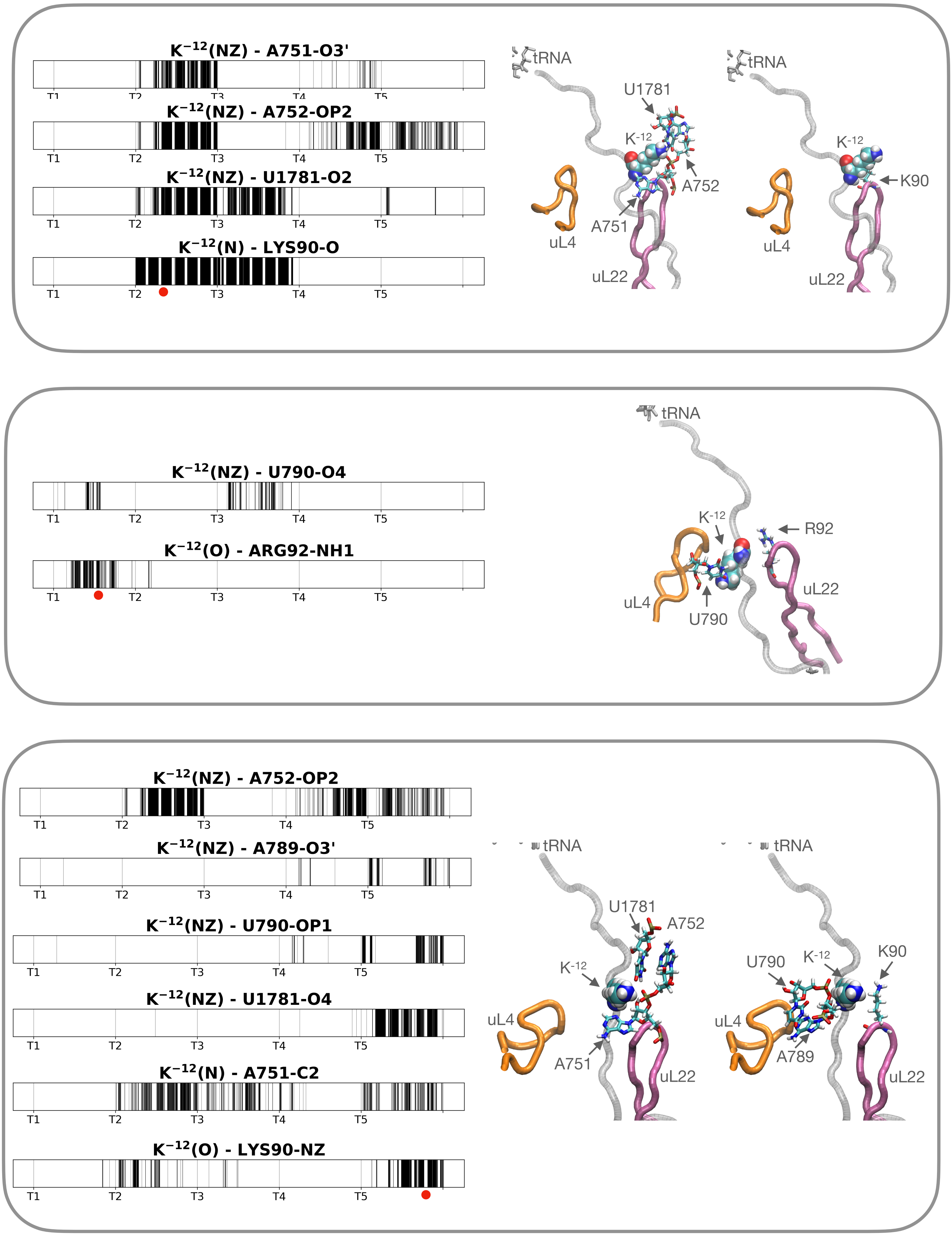


Figure S7

### Distance < 0.4 nm to N$^{-12}$

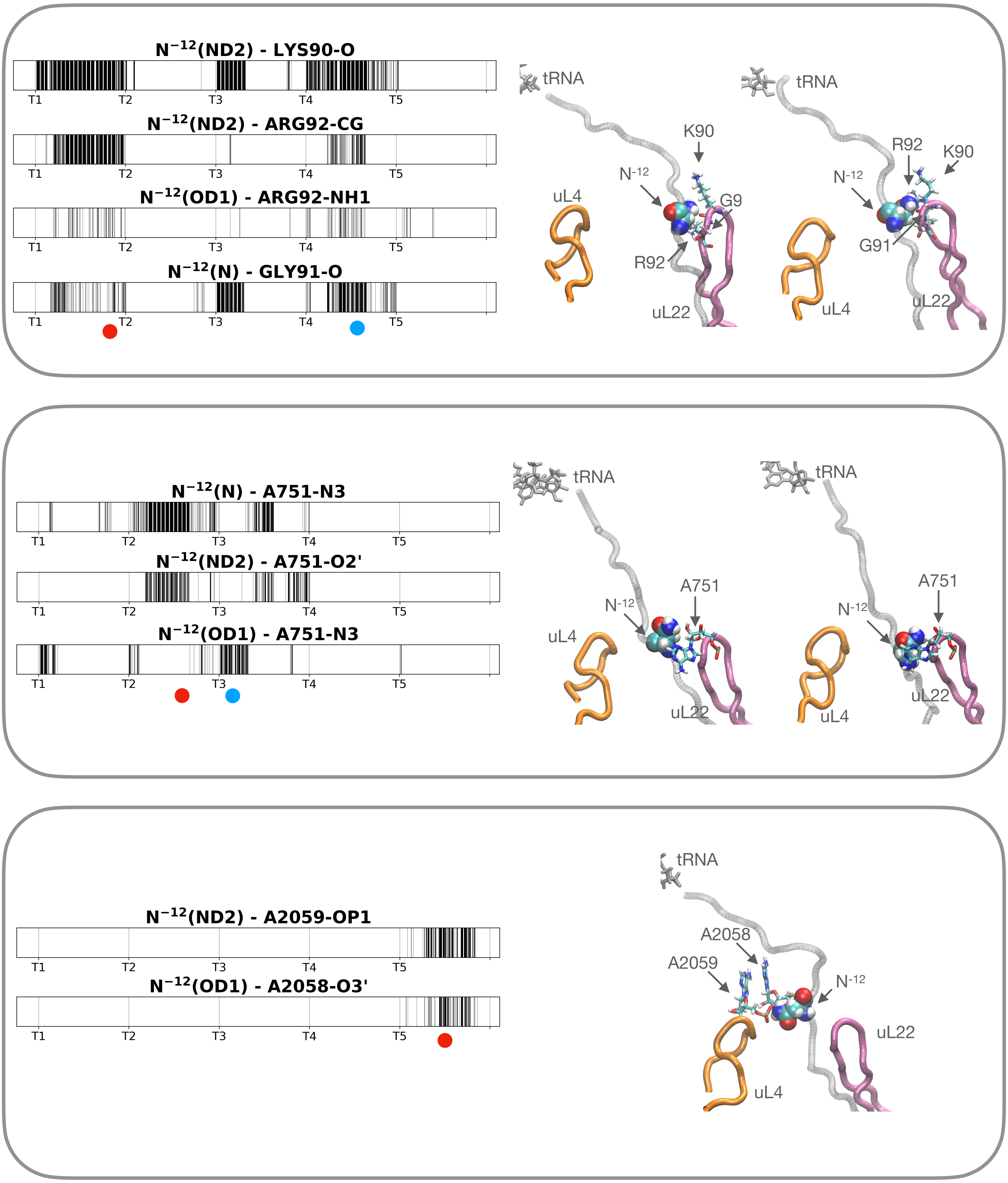


Figure S8

### Distance < 0.4 nm to $G^{-12}$

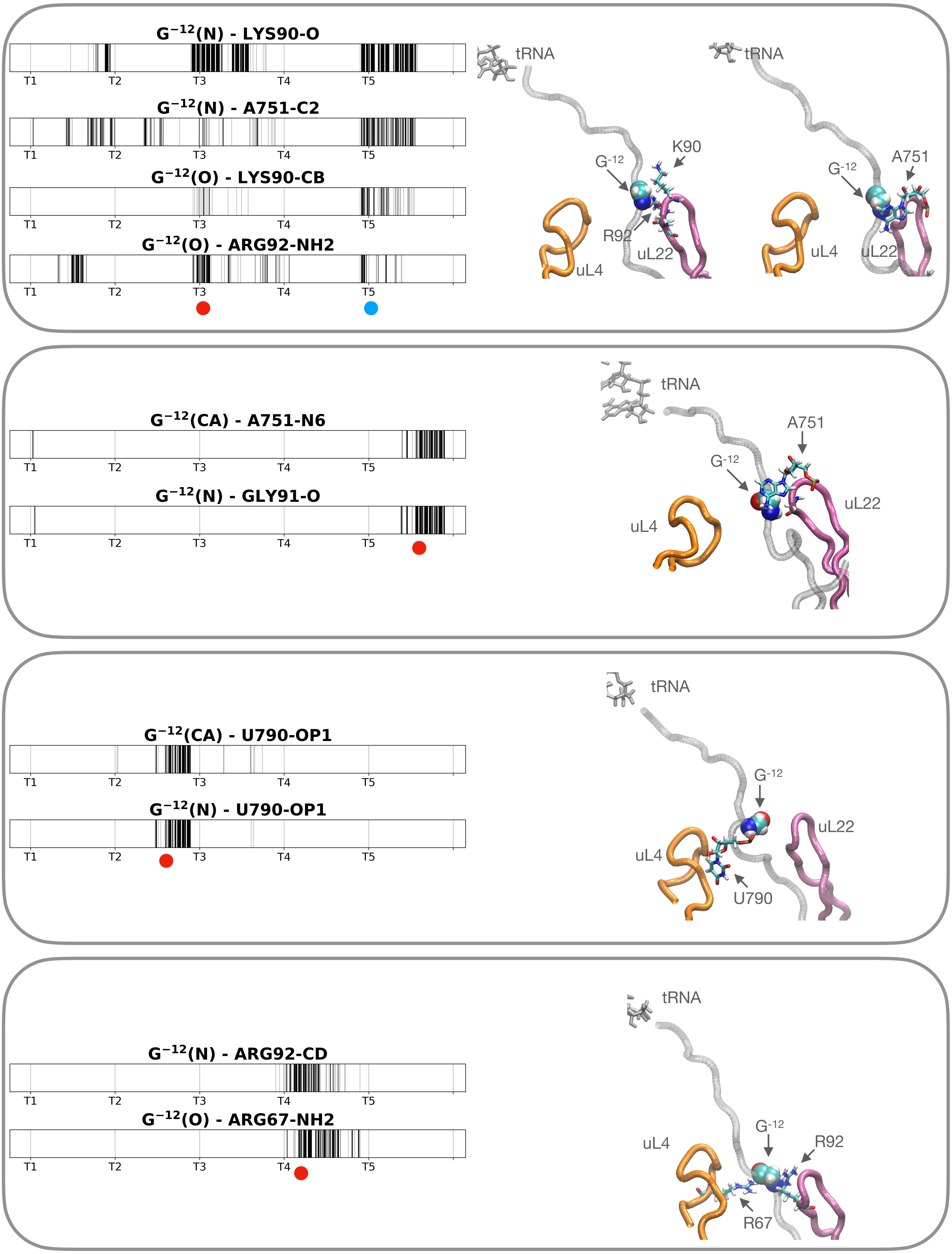


Figure S9

### Distance < 0.4 nm to $G^{-12}$ (Control)

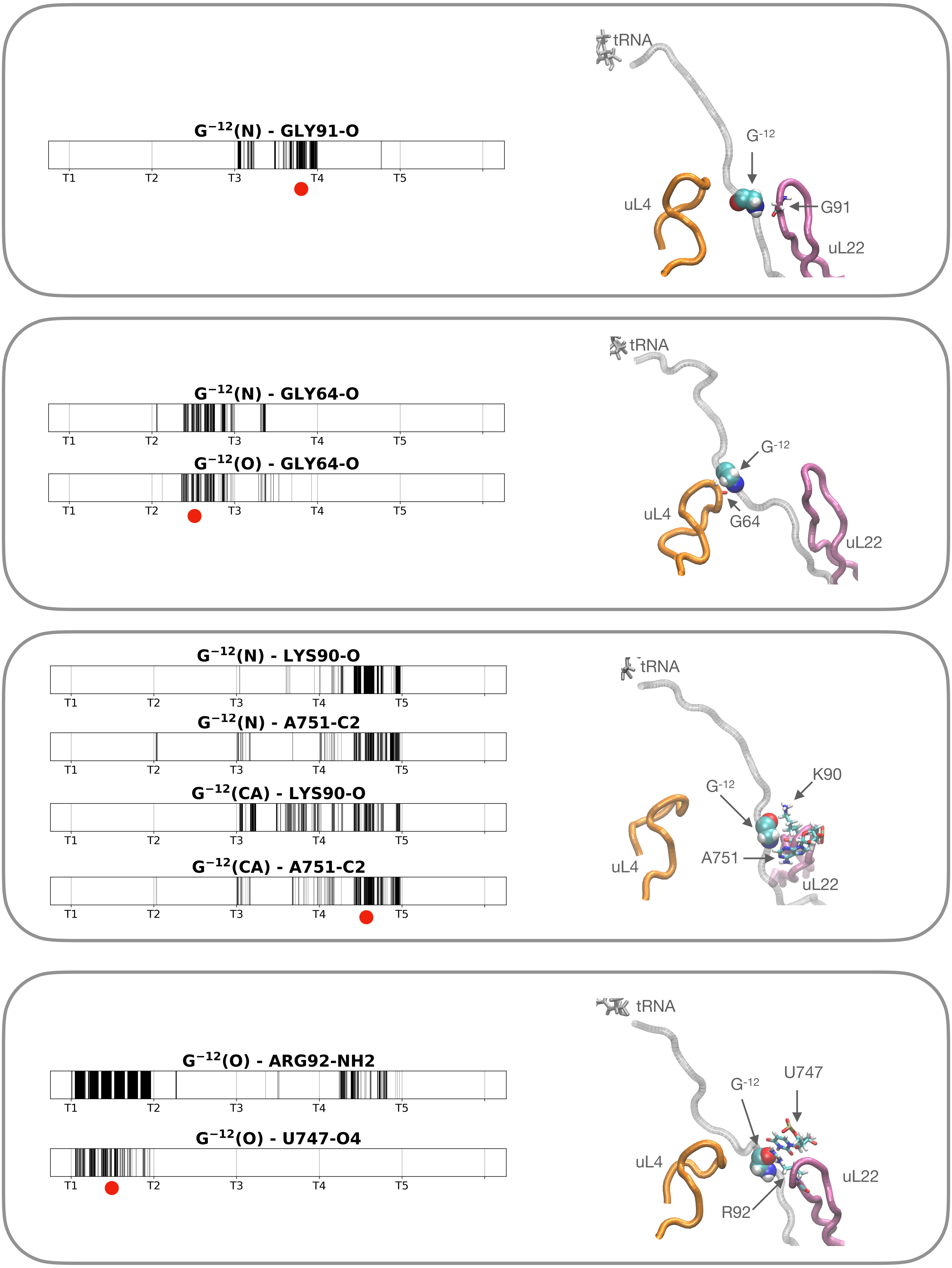


Figure S10

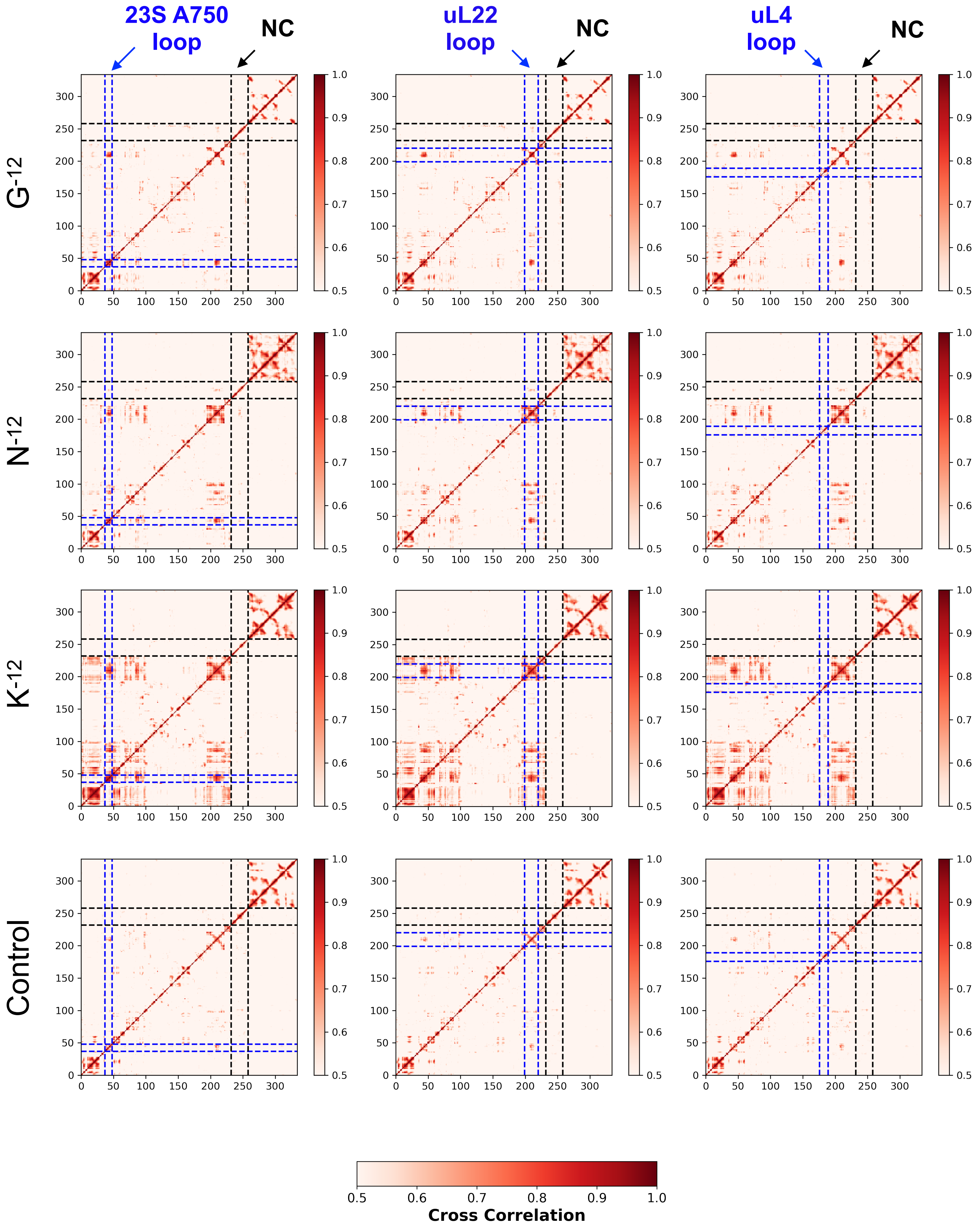
23S A750 loop
NC
uL22 loop
NC
uL4 loop
NC
G-12
N-12
K-12
Control
Cross Correlation

Figure S11

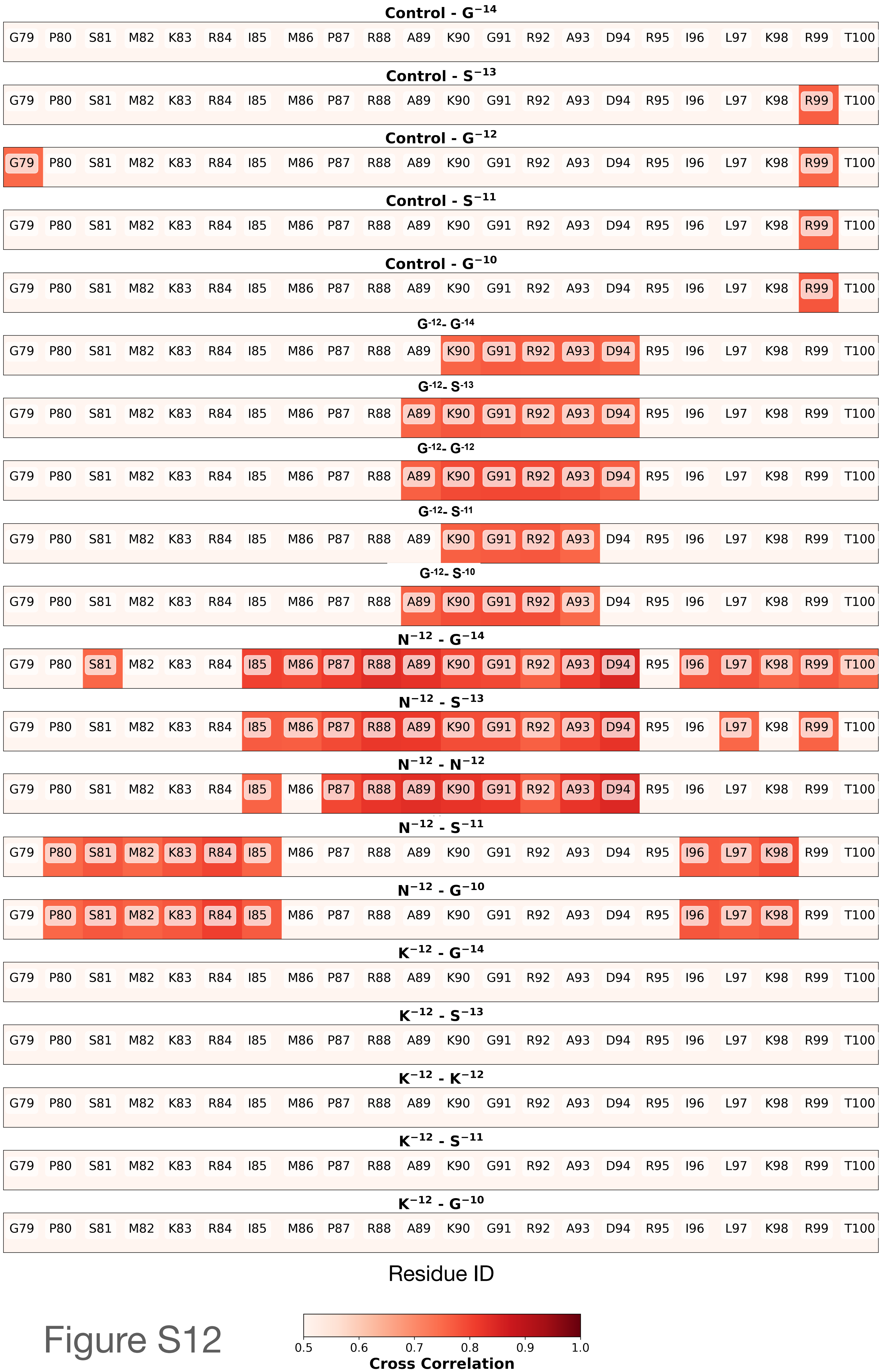

Control - G$^{-14}$
G79 P80 S81 M82 K83 R84 I85 M86 P87 R88 A89 K90 G91 R92 A93 D94 R95 I96 L97 K98 R99 T100
Control - S$^{-13}$
G79 P80 S81 M82 K83 R84 I85 M86 P87 R88 A89 K90 G91 R92 A93 D94 R95 I96 L97 K98 R99 T100
Control - G$^{-12}$
G79 P80 S81 M82 K83 R84 I85 M86 P87 R88 A89 K90 G91 R92 A93 D94 R95 I96 L97 K98 R99 T100
Control - S$^{-11}$
G79 P80 S81 M82 K83 R84 I85 M86 P87 R88 A89 K90 G91 R92 A93 D94 R95 I96 L97 K98 R99 T100
Control - G$^{-10}$
G79 P80 S81 M82 K83 R84 I85 M86 P87 R88 A89 K90 G91 R92 A93 D94 R95 I96 L97 K98 R99 T100
G$^{-12}$- G$^{-14}$
G79 P80 S81 M82 K83 R84 I85 M86 P87 R88 A89 K90 G91 R92 A93 D94 R95 I96 L97 K98 R99 T100
G$^{-12}$- S$^{-13}$
G79 P80 S81 M82 K83 R84 I85 M86 P87 R88 A89 K90 G91 R92 A93 D94 R95 I96 L97 K98 R99 T100
G$^{-12}$- G$^{-12}$
G79 P80 S81 M82 K83 R84 I85 M86 P87 R88 A89 K90 G91 R92 A93 D94 R95 I96 L97 K98 R99 T100
G$^{-12}$- S$^{-11}$
G79 P80 S81 M82 K83 R84 I85 M86 P87 R88 A89 K90 G91 R92 A93 D94 R95 I96 L97 K98 R99 T100
G$^{-12}$- S$^{-10}$
G79 P80 S81 M82 K83 R84 I85 M86 P87 R88 A89 K90 G91 R92 A93 D94 R95 I96 L97 K98 R99 T100
N$^{-12}$ - G$^{-14}$
G79 P80 S81 M82 K83 R84 I85 M86 P87 R88 A89 K90 G91 R92 A93 D94 R95 I96 L97 K98 R99 T100
N$^{-12}$ - S$^{-13}$
G79 P80 S81 M82 K83 R84 I85 M86 P87 R88 A89 K90 G91 R92 A93 D94 R95 I96 L97 K98 R99 T100
N$^{-12}$ - N$^{-12}$
G79 P80 S81 M82 K83 R84 I85 M86 P87 R88 A89 K90 G91 R92 A93 D94 R95 I96 L97 K98 R99 T100
N$^{-12}$ - S$^{-11}$
G79 P80 S81 M82 K83 R84 I85 M86 P87 R88 A89 K90 G91 R92 A93 D94 R95 I96 L97 K98 R99 T100
N$^{-12}$ - G$^{-10}$
G79 P80 S81 M82 K83 R84 I85 M86 P87 R88 A89 K90 G91 R92 A93 D94 R95 I96 L97 K98 R99 T100
K$^{-12}$ - G$^{-14}$
G79 P80 S81 M82 K83 R84 I85 M86 P87 R88 A89 K90 G91 R92 A93 D94 R95 I96 L97 K98 R99 T100
K$^{-12}$ - S$^{-13}$
G79 P80 S81 M82 K83 R84 I85 M86 P87 R88 A89 K90 G91 R92 A93 D94 R95 I96 L97 K98 R99 T100
K$^{-12}$ - K$^{-12}$
G79 P80 S81 M82 K83 R84 I85 M86 P87 R88 A89 K90 G91 R92 A93 D94 R95 I96 L97 K98 R99 T100
K$^{-12}$ - S$^{-11}$
G79 P80 S81 M82 K83 R84 I85 M86 P87 R88 A89 K90 G91 R92 A93 D94 R95 I96 L97 K98 R99 T100
K$^{-12}$ - G$^{-10}$
G79 P80 S81 M82 K83 R84 I85 M86 P87 R88 A89 K90 G91 R92 A93 D94 R95 I96 L97 K98 R99 T100
Residue ID
0.5 0.6 0.7 0.8 0.9 1.0
Cross Correlation


Figure S12

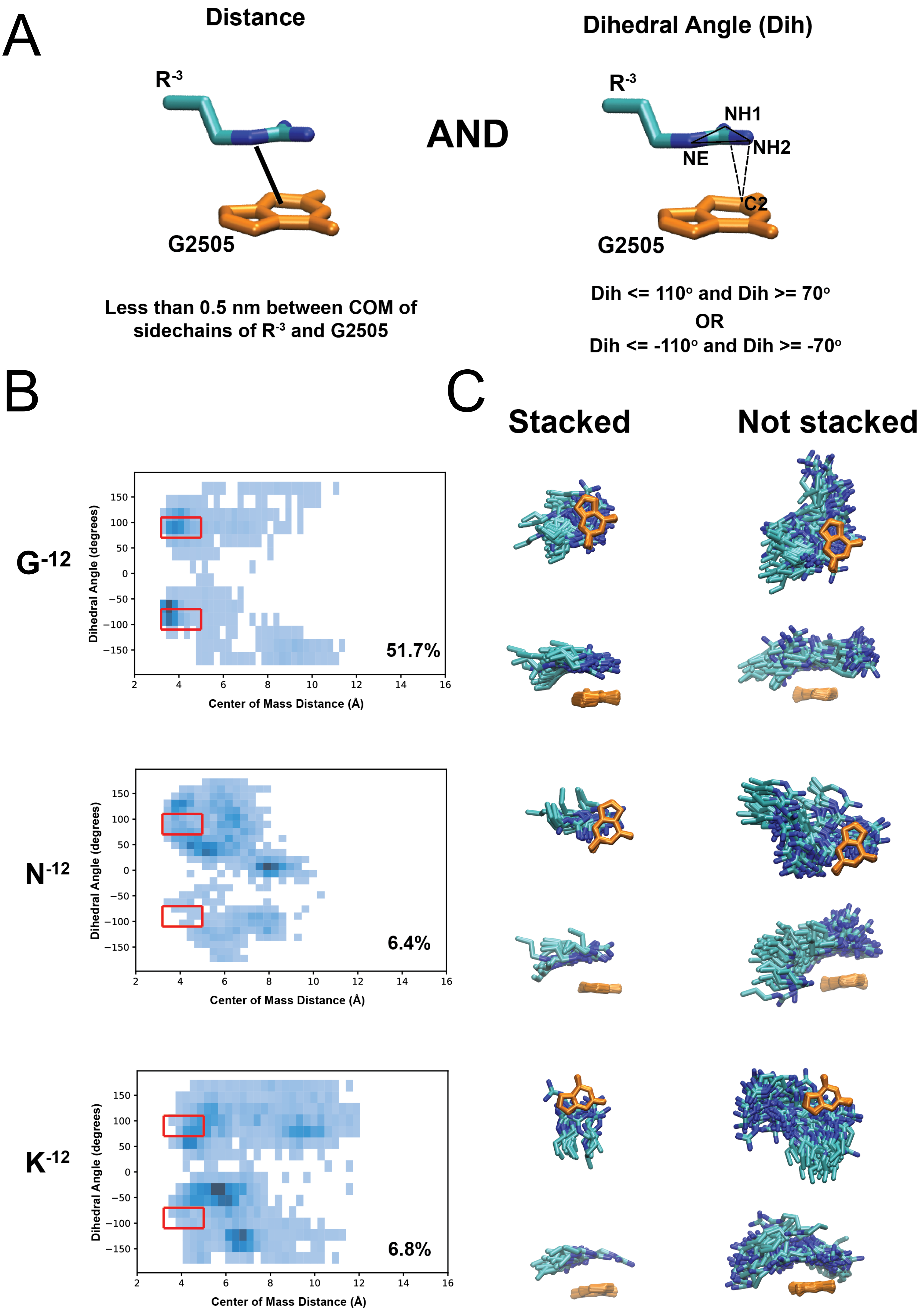
A
Distance
R-3
G2505
Less than 0.5 nm between COM of sidechains of R-3 and G2505
AND
Dihedral Angle (Dih)
R-3
NH1
NH2
NE
C2
G2505
Dih <= 110° and Dih >= 70°
OR
Dih <= -110° and Dih >= -70°
B
G-12
N-12
K-12
Dihedral Angle (degrees)
Center of Mass Distance (Å)
51.7%
6.4%
6.8%
C
Stacked
Not stacked

Figure S13
